# PfHDAC1 is an essential regulator of parasite asexual growth with its altered genomic occupancy and activity associated with artemisinin drug resistance in *Plasmodium falciparum*

**DOI:** 10.1101/2022.05.01.490232

**Authors:** Abhishek Kanyal, Heledd Davies, Bhagyashree Deshmukh, Dilsha Farheen, Moritz Treeck, Krishanpal Karmodiya

## Abstract

*Plasmodium falciparum* is a deadly protozoan parasite and the causative agent of malaria, which accounts for close to 200 million cases and 400,000 deaths every year. It has been identified to possess a tightly regulated gene expression profile that is integrally linked to its timely development during the intraerythrocytic stage. Epigenetic modifiers of the histone acetylation code have been identified as key regulators of the parasite’s transcriptome. In this study, we characterize the solitary class I histone deacetylase PfHDAC1 and demonstrate that phosphorylation is required for its catalytic activity. PfHDAC1 binds to and regulates parasite genes responsible for housekeeping and stress-responsive functions. We show that PfHDAC1 activity in parasites is crucial for normal intraerythrocytic development and that its cellular abundance is correlated with parasitemia progression. We further show that PfHDAC1 has differential abundance and genomic occupancy in artemisinin drug-resistant vs sensitive parasites and that inhibition of its deacetylase activity can modulate the sensitivity of parasites to the drug. We also identify that artemisinin exposure can interfere with PfHDAC1 phosphorylation and its genomic occupancy. Collectively, our results demonstrate PfHDAC1 to be an important regulator of basic biological functions in parasites while also deterministic of responses to environmental stresses such as antimalarial drugs.

## Introduction

Malaria is a deadly disease that affects approximately 25 million people around the world and closes in on 400,000 deaths annually, mostly in underprivileged developing countries [1]. Despite decades worth of global investment in controlling and treating the disease, it persists to ravage our well-being, especially among the vulnerable age group under 5 years old and pregnant women [1]. Among the five species that infect humans, *Plasmodium falciparum* is associated with maximum morbid manifestations and takes the lead in death tolls by the parasite [2]. Over the years, a plethora of therapeutic agents have been utilized with initial success to curb this disease, but the parasite has shown resilience and emerged resistant, to varying degrees, to most of these pharmaceutical drugs [3]. Successive utilization and failure of therapy has resulted in the selection of multidrug resistant strains of the parasite across the globe [4]. Artemisinin-based combination therapy (ACT), despite being tremendously effective in clearing parasite load, is threatened in the face of emerging artemisinin resistance across the world [5]. Artemisinin acts by eliciting a free radical cascade in parasites and compromising the functions of biomolecules by forming adducts [6]. However, *Plasmodium falciparum*, bolstered by mutations in key genes (including PfKelch13 and PfCoronin), has gradually emerged resistant to artemisinin, especially in the Greater Mekong Subregion of Southeast Asia [7, 8]. Recent studies have linked artemisinin resistance to changes in lipid metabolism and an overall reduction in heme endocytosis and processing [9, 10]. Comparative multiomics profiling of artemisinin-resistant parasites has revealed substantial reconfiguration of gene expression networks in response to the drug to reduce its activation and enhance the mitigatory homeostatic functions in parasites [11, 12]. This highlights the need to uncover these regulatory and adaptive mechanisms in the parasite.

*Plasmodium falciparum* displays immense genetic and phenotypic plasticity, allowing it to mutate and adapt to environmental pressures. Furthermore, despite being placed very early in the evolutionary tree of life, the parasite has significantly invested in adequate utilization of its genome [13]. The parasite displays strictly coordinated expression of key genes over the 48-hour intraerythrocytic development cycle, where it transitions through phases of increasing metabolic activity and replication, culminating in optimal utilization of host nutrients and then egressing to infect a fresh set of cells [14, 15]. Characterized by relatively AT-rich genomes, multiple *Plasmodium* species have come to utilize a host of epigenetic mechanisms (especially histone variants and modifications) to orchestrate gene expression regulation [16]. Modifiers of the histone acetylation code in *P. falciparum* play a crucial role in basal transcriptional regulation as well as response to abiotic (fever cycles) and biotic stresses (immune evasion) [17]. Due to their sequence proximity to counterpart molecules in earlier eukaryotes, histone acetyltransferases and deacetylases have been heavily investigated as potential therapeutic targets for epigenetic modulators and inhibitors. Interestingly, HDACs have emerged as potential epigenetic regulators of molecules that have direct implications in determining the efficacy of various therapeutic drugs (especially chemotherapy). Pharmacological inhibition and genetic depletion of HDACs have been associated with deregulated expression of transporter molecules such as MDR1 (multidrug resistance 1) and BCRP (breast cancer resistance protein), allowing for the emergence of resistance to chemotherapy [18]. This becomes a fact of relevance and concern since histone deacetylase inhibitors adapted from anti-cancer drug panels using a piggy-back approach have shown great promise with their anti-*Plasmodium* activity [19]. Despite this success, the targets of these compounds are relatively poorly understood. There are 3 major classes of histone deacetylases in *Plasmodium falciparum*: Zn2+ cofactor-dependent class I (PfHDAC1) and II (PfHda1 and PfHdaII) and NADH cofactor-dependent class III/sirtuins (PfSir2a and PfSir2b) [17]. PfHDAC1 is the ‘lone-wolf’ of the Class I HDACs and is still very poorly studied in *P. falciparum*. Genome-wide mutagenesis screens have suggested the gene to be essential in both *P. falciparum* and *P. berghei* [20, 21]. This underlines the potentially critical role HDAC1 plays in *Plasmodium* biology. Numerous drugs (including entinostat (MS-275), romidepsin, valproic acid, LMK-35 and NSAID with anti-parasite activity are believed to operate by targeting PfHDAC1 [19, 22-25]. *Plasmodium falciparum* is currently treated by strictly regulated regimens of artemisinin-based combination therapies (ACT). However, in the last decade, there have been breakthrough reports of the emergence and spread of artemisinin resistance in the field, especially in Southeast Asia. A population transcriptomics study conducted on artemisinin-resistant and -sensitive parasites from Southeast Asia by Mok et al. identified PfHDAC1 as one of the most significantly downregulated genes (among the bottom 5% of the genes arranged by their z-score) [26]. Given the evidence of the role of HDAC inhibition in the emergence of chemotherapeutic resistance in mammalian systems, it becomes highly interesting to cross-examine the role of PfHDAC1 in governing resistance to antimalarial compounds (chiefly artemisinin) against *P. falciparum*.

In this study, we perform a basic characterization of PfHDAC1, especially in the context of its gene regulatory role. HDAC1 in higher-order eukaryotes has been associated with regulating key biological functions in tandem with chromatin remodelling associated with regulation of the cell cycle, developmental fates, response to therapeutic pressure, DNA replication/damage repair, autophagy, and protein quality control [27-32]. Aberrant expression of HDAC1 in mammalian cells has been associated with neoplastic trends as well as resistance to drug therapy leading to failure of treatment regimens [33]. This puts HDAC1 at the nexus of basal as well as stress-responsive roles in organisms. We aimed to profile the deacetylase activity of PfHDAC1, its regulation by posttranslational modification, identification of its gene targets and the biological functions that it governs in the parasite. We further investigated the effect of alteration of PfHDAC1 activity on the sensitivity of parasites to artemisinin and its potential role in regulating gene expression leading to the emergence of artemisinin resistance.

## Results

### Recombinant PfHDAC1 interacts with the kinase PfCKII-α, esulting in its phosphorylation and catalytic activation

Although PfHDAC1 has been described as a target of multiple highly potent inhibitors, a thorough genetic and biological characterization of PfHDAC1 function is lacking in *P. falciparum* [19, 34-36]. To characterize its enzymatic activity, we cloned, expressed, and purified histidine- and GST-tagged PfHDAC1 (Figure 1A and 1B). Previous studies provide a strong indication that phosphorylation by casein kinase (CKII-α) is required for the deacetylase activity of HDAC1 in mammalian systems [37]. A survey of the existing blood stage total and phospho-proteome literature indicates that PfHDAC1 is phosphorylated at serine residues, specifically S391, S397 and S440 [38, 39]. We analyzed the PfHDAC1 amino acid sequence on the NetPhos-3.1 web tool, which predicts/scores the potential phosphorylation sites on protein sequences and the associated kinase enzymes [40]. The tool predicted a strong phosphorylation potential for the S391, S397 and S440 residues and phosphorylation by the CKII kinase at these sites (Supplementary Fig. 1A-C). We thus sought to test whether PfCKII (more specifically PfCKII-α) could be responsible for the phosphorylation of PfHDAC1. First, we generated an α-PfHDAC1 antibody against the full-length protein (Fig 1C) and confirmed its specificity by immunoprecipitation followed by mass spectrometry on parasite lysate. PfHDAC1 was identified in the mass spectrometry analysis in PfHDAC1 immunoprecipitation, whereas it was absent in the IgG immunoprecipitation. We also cloned, expressed, and purified PfCKII-α-GST to homogeneity (Supp Fig. 2A and 2B). To further validate whether the two proteins PfHDAC1 and PfCKII-α interact *in vitro*, we incubated the purified recombinant proteins together followed by immunoprecipitation using our PfHDAC1 antibody. An IgG pulldown was used as a control. The immunoprecipitated fractions were resolved on a 10% SDS-PAGE gel, and western blotting was performed using an α-GST antibody to probe for coimmunoprecipitated PfCKII-α. We detected an enrichment of PfCKII-α in the PfHDAC1 IP but not in the IgG IP (Supp Fig. 2C), confirming the *in vitro* interaction of PfHDAC1 and PfCKII-α.

**Figure 1:**
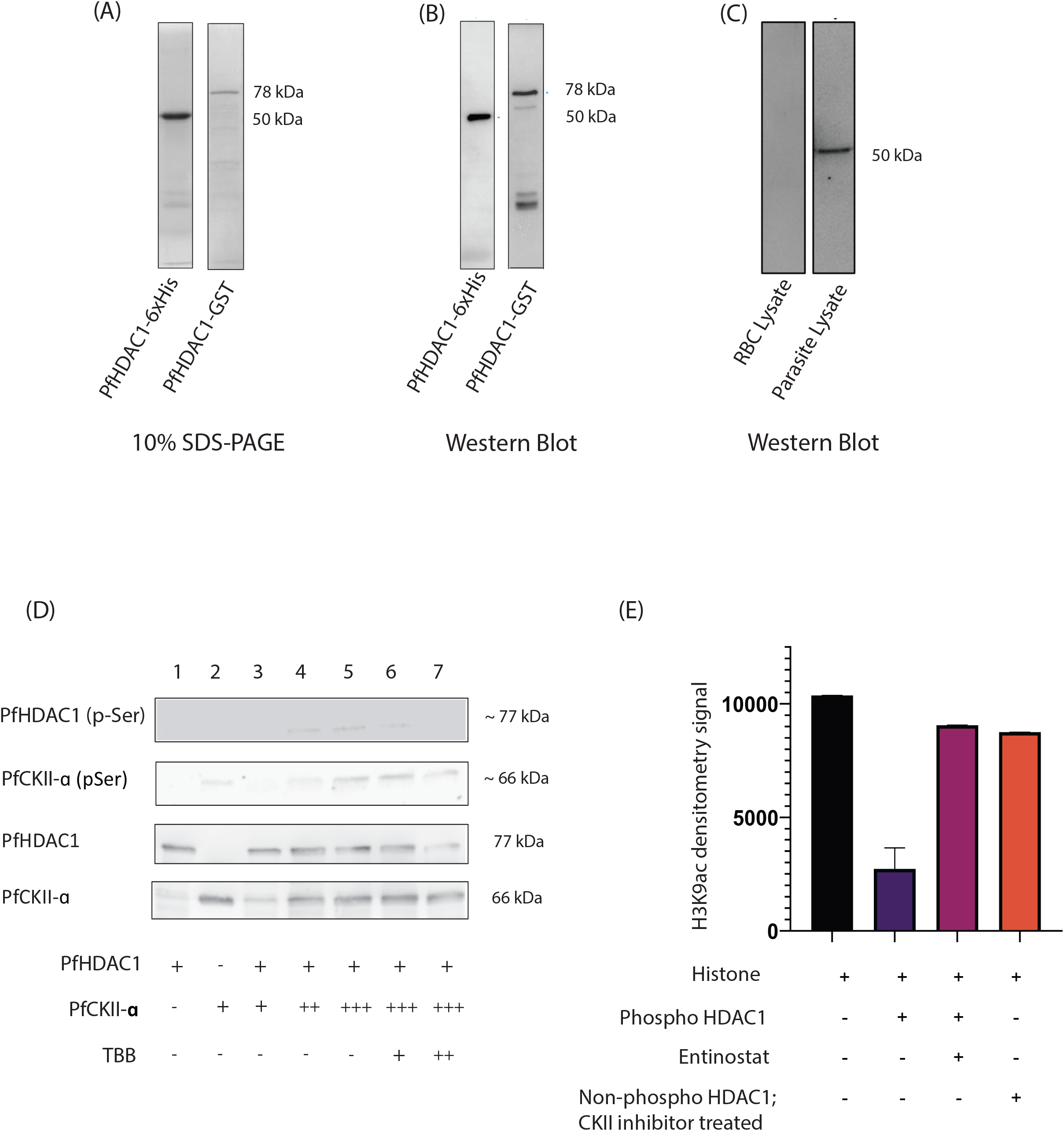
PfCKII-α-mediated phosphorylation of PfHDAC1 is necessary for its enzymatic activity. (A) SDS-PAGE (10% resolving) and (B) Western blotting for purified recombinant PfHDAC1-GST and PfHDAC1-6xHis. (C) Western blotting for parasite lysate and uninfected RBC control lysate probed with purified in-house PfHDAC1 antibodies. (D) Western blot (using α-phospho Ser/Thr antibody in top two panels and α-GST antibody bottom two panels) for in vitro kinase activity assay using purified recombinant PfHDAC1 and PfCKII-α. The reaction mix components are as specified in the following table. (E) Histogram representing the western blot signal for H3K9ac probed in the in vitro deacetylase activity assay. The reaction components are specified in the following table.

We next sought to validate whether this interaction was functionally relevant and resulted in the phosphorylation of PfHDAC1. We set up an *in vitro* kinase activity assay in which recombinant PfHDAC1 was incubated with increasing amounts of PfCKII-α (Fig. 1D). We also set up two controls where the kinase reaction was spiked with increasing doses of the PfCKII-α-specific inhibitor TBB. The incubated mixes were run through 10% SDS-PAGE and subsequently probed with an α-phosphoserine/threonine antibody to recognize any phosphorylation-modified forms of PfHDAC1 or PfCKII-α (Fig. 1D). We observed an appearance of the phosphorylated form of PfHDAC1 in reactions with PfCKII-α coincubation. The control reaction without PfCKII-α did not show any band for phospho-PfHDAC1. In the reaction mixes incubated with the PfCKII-α inhibitor, we observed inhibition of the phosphorylated PfHDAC1 band. We also noticed that PfCKII-α has autophosphorylation activity, which is also inhibited in the presence of TBB. These results indicate that PfHDAC1 and PfCKII-α can interact *in vitro* and that this interaction results in the phosphorylation of PfHDAC1.

To investigate whether the phosphorylation of PfHDAC1 could contribute to its catalytic activity, we performed *in vitro* histone deacetylase assays using histones isolated from *P. falciparum* as substrates and PfCKII-α-treated PfHDAC1 (phosphorylated) as the enzyme (Fig. 1E and Supp Fig. 2D). Phosphorylated PfHDAC1 displayed histone deacetylase activity, as we observed depletion of H3K9ac histone modification on the substrate, which was confirmed by western blotting. Furthermore, we used the class I-specific histone deacetylase inhibitor entinostat (MS-275) and observed that it suppresses the deacetylase activity associated with phosphorylated PfHDAC1. Additionally, a control reaction with nonphosphorylated recombinant PfHDAC1 (from TBB inhibited PfCKII-α kinase reaction on PfHDAC1) resulted in no deacetylase activity. Collectively, our results suggest that PfCKII-α-dependent phosphorylation of PfHDAC1 is important for robust deacetylase activity, which can be inhibited by a class I-specific histone deacetylase inhibitor.

### PfHDAC1 genomic occupancy is associated with a wide range of housekeeping and stress-responsive genes

To identify the genome-wide targets of PfHDAC1 with chromatin immunoprecipitation followed by sequencing (ChIP-seq), we generated a transgenic parasite line in which endogenous PfHDAC1 was tagged with 2xFKBP-GFP at the C-terminus (Fig. 2A and 2B) (adapted from [41]). We validated the GFP tagging of PfHDAC1 using the α-GFP antibody by western blotting (Fig. 2C) and GFP fluorescence using confocal microscopy (Fig. 2D). To identify the targets of PfHDAC1, we performed ChIP sequencing using anti-GFP antibodies in two replicates. We chose early trophozoite stage parasites at approximately 24 hours post invasion (HPI) for ChIP-seq since this stage is characterized by a surge in parasite transcriptome as well as robust PfHDAC1 expression. Through ChIP-seq, we identified 1409 genes with >=2-fold enrichment across two replicates for PfHDAC1 in *P. falciparum* (Fig. 2E, 2F and Supplementary Fig. 2E). The PfHDAC1 peaks were enriched predominantly around the transcription start/end sites of target genes with coverage extending into the 5’ and 3’ UTRs. We would like to point out that with version 53 of the *P. falciparum* genome, many genes (but not all) have now been assigned 5’ and 3’ UTR information. Thus, for PfHDAC1 target genes with designated 5’/3’UTR information, the start/stop sites were transcriptional, and for those without this information, the start site used as a reference was translational. Only a small number of coding sequences (CDSs) were identified to be bound by PfHDAC1 (Fig. 2E). Gene Ontology enrichment analysis of the target genes revealed biological functions associated with movement within the host cell, phosphorylation, cell signalling, haemoglobin metabolism, redox homeostasis, protein folding, regulation of splicing, autophagosome assembly, and cell cycle to be linked with PfHDAC1 (Fig. 2G). Thus, we found PfHDAC1 gene targets to be associated with both housekeeping (entry into host cell, hemoglobin metabolism, cell cycle) and parasite stress responsive functions (protein folding, redox homeostasis). For a functional validation of the target genes, we chose to inhibit PfHDAC1 with Class I HDAC-specific inhibitors (romidepsin and entinostat) and checked the expression of several PfHDAC1 ChIP target genes via RT-qPCR. Few genes belonging to the protein refolding, protein ubiquitination and redox response pathways were investigated in this way. The expression of the chaperones heat shock protein 70 (PfHSP70) and PfBiP and the redox response proteins ER-oxidoreductase and ubiquitination-associated PfPEX-E2 and PfRBX-E3 was found to be highly upregulated upon PfHDAC1 inhibitor treatment (Fig. 2H). Thus, we identified that PfHDAC1 has genomic occupancy over genes associated with housekeeping functions and stress responses.

**Figure 2:**
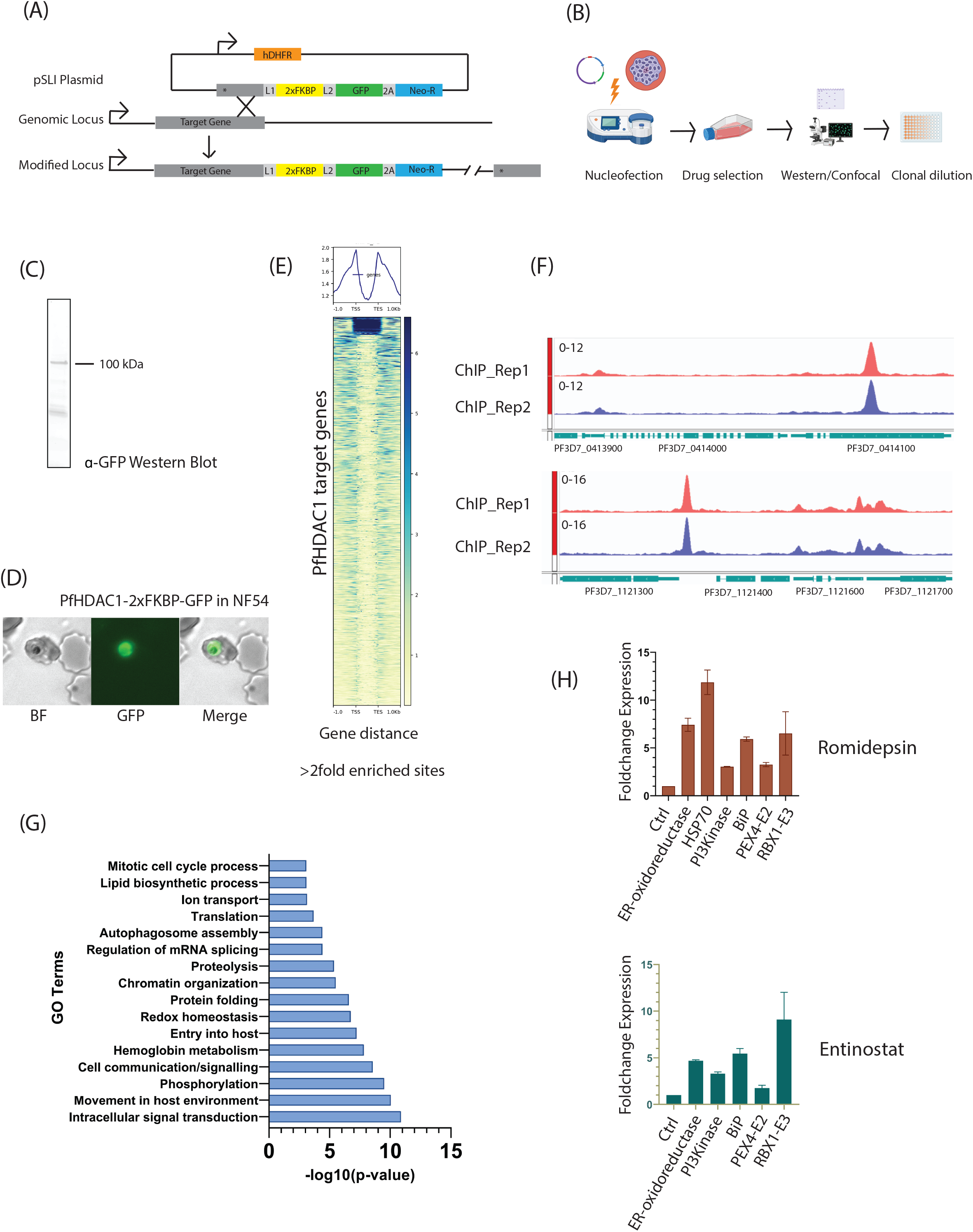
PfHDAC1 binds to *Plasmodium* genes associated with house-keeping and stress-responsive functions. (A) Schematic of the PfHDAC1-2xFKBP-GFP knockin construct and recombination/integration into the genomic locus. (B) Schematic of transfection, drug selection and clonal selection of the transgenic PfHDAC1-2xFKBP-GFP parasite line. (C) Western blotting (α-GFP) and (D) confocal microscopy (α-GFP) to validate the knockin line. (E) Heatmaps displaying the genomic enrichment of PfHDAC1 at >= 2-fold enriched target genomic loci. (F) IGV browser sample shots of tracks displaying enrichment peaks of PfHDAC1. (G) Histogram representing the Gene Ontology enrichment of biological processes under PfHDAC1 occupancy. (H) Histogram representing the upregulation of selected PfHDAC1 target genes upon romidepsin and entinostat treatment (qRT-PCR data).

### PfHDAC1 is a regulator of parasite intraerythrocytic development

Our initial assessment of PfHDAC1 gene targets revealed enrichment of several important biological pathways, including host cell entry/egress, cellular signaling, hemoglobin metabolism and the cell cycle. Given that these are processes associated with parasite development and the progression of infection in host cells, we decided to observe the effects of PfHDAC1 inhibition on intra-erythrocytic development and re-invasion into fresh RBCs. Class I HDACs have been well characterized for their role in cell cycle regulation [32]. HDAC1 is known for regulating the expression of cyclins and cyclin-dependent kinases and is an integral factor governing cell cycle checkpoint dynamics in mammals [42]. We inhibited PfHDAC1 using a continuous sublethal dosage of romidepsin (a class I HDAC inhibitor) and followed intraerythrocytic cell cycle progression via microscopic examination at 8-hour intervals (starting at 6-hour post-invasion mark) (Fig. 3A and 3B). Since the cell cycle progression in *P. falciparum* is also linked with progressive DNA replication in the process of erythrocytic schizogony, we estimated the DNA content changes through SYBR fluorescence readout at each of these timepoints (Fig 3C). We counted the individual stages (ring/trophozoite/schizont) present at each timepoint and estimated the stage progression (or inhibition) of parasites over time (Fig 3B). At 22 hours post invasion, the mock-treated parasites had almost completely transitioned from the ring to the trophozoite stage, whereas a small proportion of the inhibitor-treated cells were still found to be lagging in the ring stage or not fully transitioned into trophozoite morphology. Trophozoite development was closely monitored in mock-treated cultures and showed gradual enlargement of the nuclear volume associated with maturation of the trophozoite (noted at 30 HPI). In the inhibitor-treated culture, a vast majority of trophozoites showed stunted maturation, and only a few of those developed into full-sized trophozoites (as referenced in mock-treated culture). This hinted at a strong inhibition of morphological development possibly associated with stunted DNA replication in PfHDAC1-inhibited parasites. Furthermore, segmented schizonts were observed at the 46 hour post-invasion time point in mock-treated parasites along with egressing parasites. In contrast, the romidepsin-treated culture showed a large proportion of immature schizonts with delayed segmentation. While a significant proportion of the mock-treated cells had undergone egress and reinvasion by 54 hours post invasion/6 hours post invasion of cycle 2, the romidepsin-treated parasites showed comparatively less reinvasion into fresh RBCs. Schizonts persisted in romidepsin-treated culture at least 6-8 hours after the second invasion cycle had already begun in the mock treatment culture, resulting in a delayed second wave of infection (Fig. 3A and 3B). We also estimated the DNA content of romidepsin-treated parasites and observed a significant drop at 16 hours post treatment compared to the control parasites (Fig. 3C). Interestingly, the DNA content gradually increased to 75% of mock-treated parasites in romidepsin treatment at 40 hours post invasion compared to the control parasites (Fig 3C). The 25% deficit of DNA fluorescence in romidepsin-treated parasites may reflect the loss of DNA replication observed upon PfHDAC1 inhibition. Interestingly, the decrease in DNA content upon romidepsin treatment was corroborated by a decrease in overall parasitaemia (11% in the control and 7% upon romidepsin treatment) in the second intraerythrocytic developmental cycle (Fig. 3D). Thus, PfHDAC1 inhibition was shown to delay cell cycle progression compounded with defects in the proper morphological development of parasites and a reduction in proliferation over the next cycle of intraerythrocytic development.

**Figure 3:**
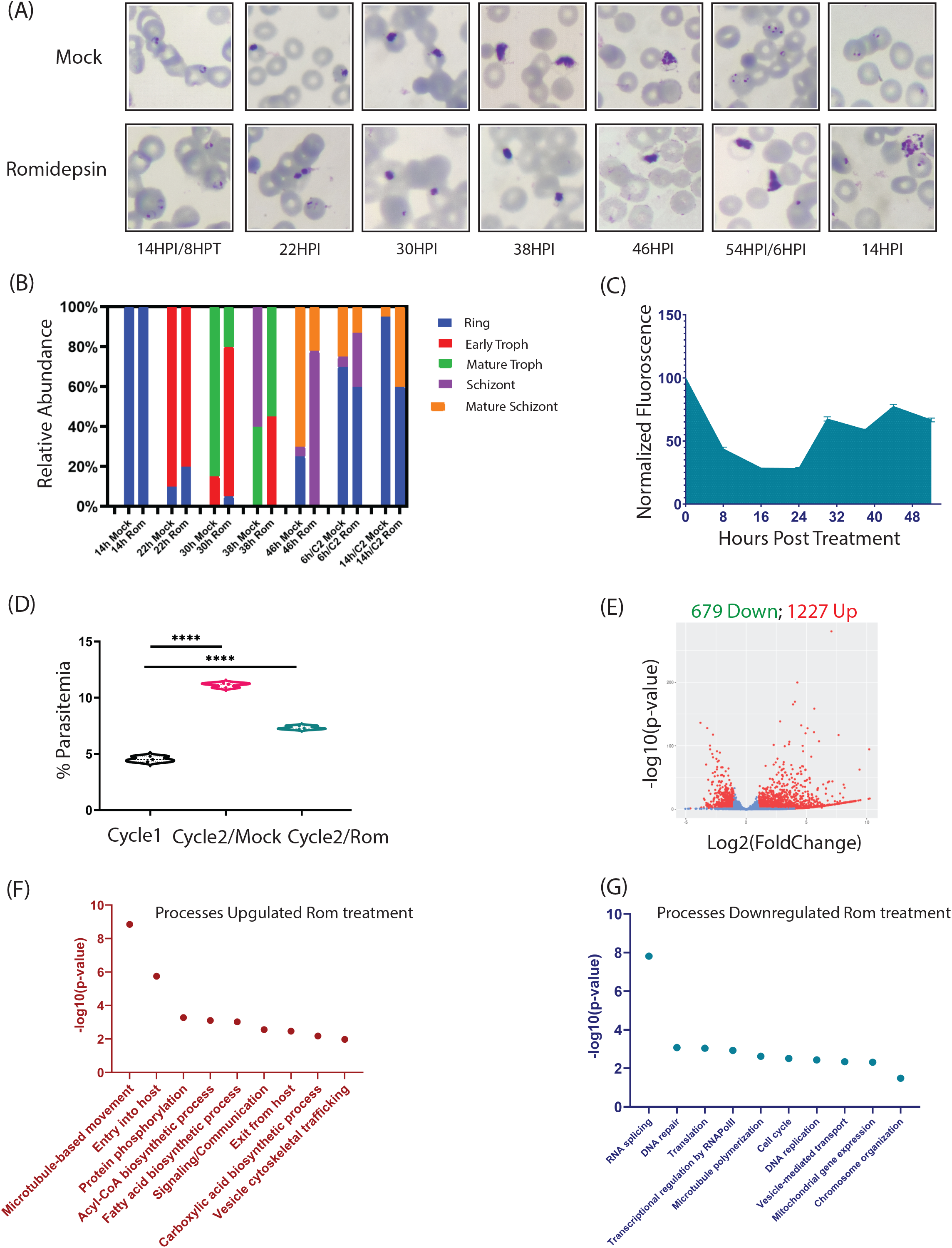
PfHDAC1 governs intraerythrocytic development, DNA replication, and infection progression in *Plasmodium falciparum*. (A) Giemsa-stained smear panel comparing parasite progression through the intraerythrocytic development cycle upon mock and romidepsin treatment. Images were taken at 8-hour intervals from 6HPI onwards. (B) Histograms representing the relative proportion of parasites from each of the three major stages (ring, trophozoite and schizont) in mock-vs romidepsin-treated parasite cultures. (C) Area plot depicting the skewed DNA content in romidepsin-treated parasites normalized to the mock control. (D) Dot plot representing the lower reinvasion rates and parasitemia progression in romidepsin-treated parasites compared to the mock control. (Statistical analysis used: Student’s unpaired t-test; **** represents p-value <0.0001) (E) Volcano plot representing the differential gene expression upon romidepsin treatment of parasite culture. Dot plots representing the biological processes (F) upregulated and (G) downregulated upon romidepsin treatment in parasites.

To identify the transcriptional changes taking place in parasites upon PfHDAC1 inhibitor treatment, we performed RNA sequencing on 21-24 HPI trophozoite stage parasites treated with romidepsin (360 nM; 2XIC_50_ for 3 hrs). We observed deregulation of 1906 genes (1227 upregulated and 679 downregulated genes; log2fold-change >= -1/1; p-value<=0.05, mean count>=10) with PfHDAC1 inhibitor treatment (Fig. 3E). Among the biological processes upregulated were entry into host, microtubule-based locomotion, fatty acid metabolism and cell signaling (Fig. 3F and Supplementary Fig. 4A). We observed the downregulation of genes associated with critical biological processes, such as RNA splicing/metabolism, transcription by RNA polymerase II, translation, DNA damage repair, cell cycle, DNA replication and vesicle-mediated transport (Fig. 3G and Supplementary Fig. 4B). We investigated the specific cell cycle- and DNA replication-associated genes that were tweaked upon PfHDAC1 inhibition. Centrin-1, putative spindle and kinetochore associated protein, anaphase promoting complex protein 1, SMC protein 1, DNA replication licensing factors MCM3/MCM7, replication factor C subunit 4/5 and DNA polymerase 1 were among the set of genes found to be downregulated upon PfHDAC1 inhibition. Taken together, our results indicate that PfHDAC1 activity in the parasite is required for the optimal expression of several genes associated with crucial biological programs linked to asexual development and host cell reinvasion of the parasite.

### PfHDAC1 overexpression is associated with a growth advantage in parasites linked with enhanced expression of host cell invasion genes

Driven by our observation of the effect of PfHDAC1 inhibitor treatment on the phenotype of parasites, we were curious to investigate the effects of increased PfHDAC1 levels in cells. We tested whether overexpression of PfHDAC1 would have converse effects on the growth and proliferation of the parasites. For this purpose, we episomally overexpressed a GFP-tagged version of PfHDAC1 (driven by the calmodulin promoter) using the pDC2-Cam-GFP overexpression plasmid. A glmS ribozyme sequence was introduced downstream of PfHDAC1-GFP (Fig. 4A), which allowed us to regulate PfHDAC1 expression levels. The overexpression lines were confirmed with α-GFP confocal microscopy and western blotting (Fig. 4B and 4C). The resultant PfHDAC1-GFP-glmS overexpression construct could be regulated at the posttranscriptional level by supplementing glucosamine ligand to the culture media, resulting in tunable overexpression of PfHDAC1 (Fig 4C). A GFP-glmS overexpression line was used as a control. Parasite growth was recorded as a fold change relative to the starting Day 0 parasitemia (at 1% starting parasitemia and 2% hematocrit) for the control (GFP-glmS) and test (PfHDAC1-GFP-glmS) parasites over a 3-cycle/6-day duration (Fig 4D). Samples were taken once every 24 hours for flow cytometry-based counting. Parasitaemia in the PfHDAC1 overexpression line was observed to increase at a faster rate than that in the GFP control line with every cycle (50% higher parasitemia at cycles 2 and 3). Thus, overexpression of PfHDAC1 appears to increase the proliferation of parasites with each cycle.

**Figure 4:**
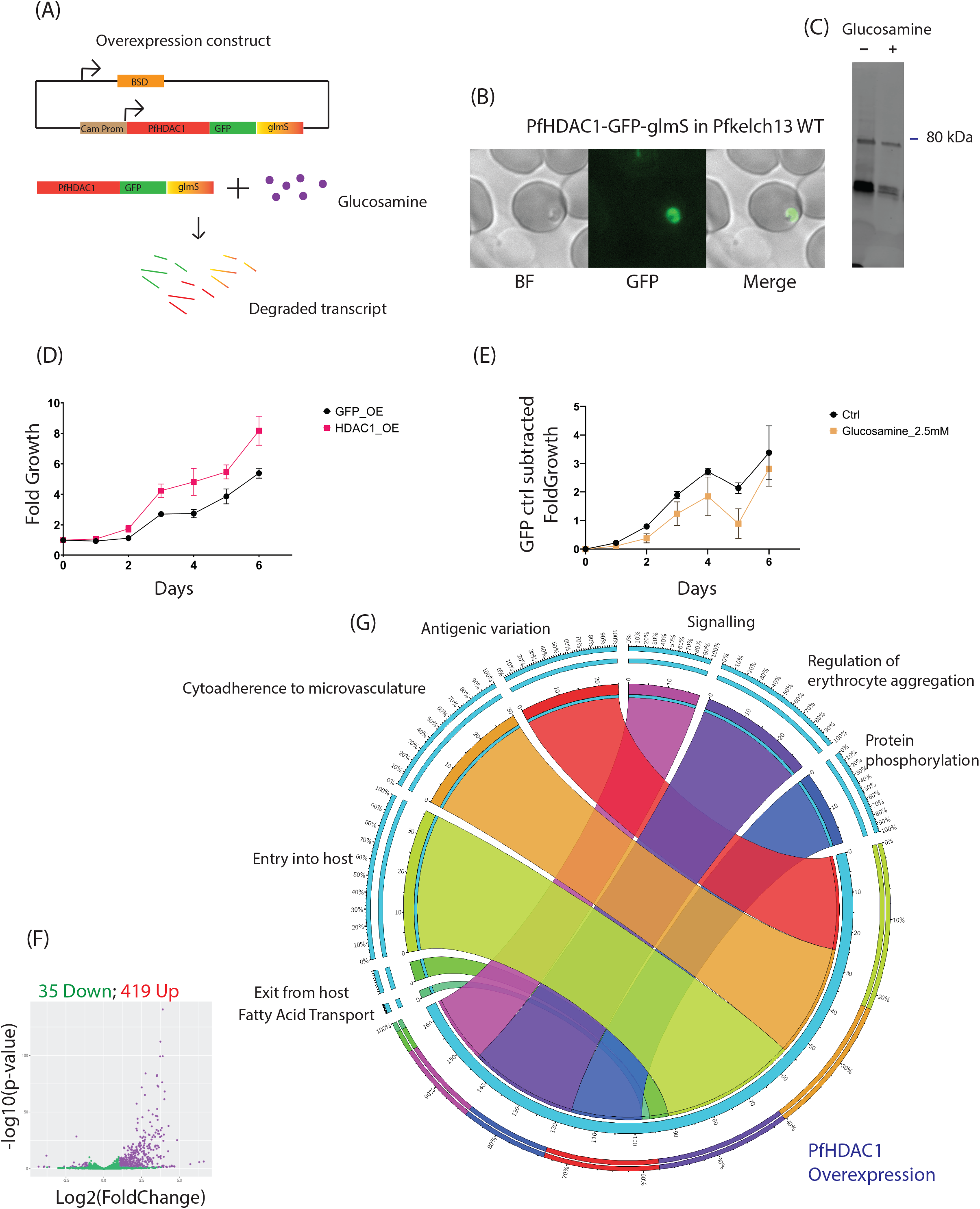
PfHDAC1 overexpression is associated with a proliferative advantage in parasites. (A) Schematic of the pDC2 plasmid used for calmodulin promoter-driven episomal overexpression of PfHDAC1-GFP-glmS. The ribozyme-tagged transcript can be degraded upon the addition of glucosamine ligand. (B) Confocal microscopy-based validation of PfHDAC1-GFP-glmS expression. (C) Western blotting-based validation of PfHDAC1-GFP-glmS overexpression in parasite lysate and its mild depletion upon glucosamine treatment. (D) Line plot demonstrating the growth advantage observed in PfHDAC1-GFP-glmS overexpression parasites compared to the control GFP-glmS overexpression line. The data is representative of PfKelch13WT genotype parasites. (E) Line plot representing the GFP-glmS subtracted growth curve for PfHDAC1-GFP-glmS line to demonstrate the drop in growth advantage observed upon 2.5 mM glucosamine treatment compared to mock treatment control. The data is representative of PfKelch13WT genotype parasites. (F) Volcano plot representing differential gene expression upon PfHDAC1 overexpression. Significantly deregulated genes are colored purple based on log2foldchange cutoff +/-1 and p-values 0.05. (G) Circos plot representing the biological processes enriched for genes upregulated with PfHDAC1 overexpression in parasites.

To test the reversibility of the phenotype induced by PfHDAC1 overexpression, we dosed the parasite culture with 2.5 mM glucosamine, which, as expected, reduced PfHDAC1-GFP-glmS levels (as validated by α-GFP western blot) (Fig 4C). Glucosamine-treated GFP-glmS parasites were used as a control. We plotted the GFP-glmS subtracted growth curves for PfHDAC1-GFP-glmS parasite lines over a 6-day growth assay (glucosamine vs mock treatment) (Fig. 4E). While mock-treated PfHDAC1 overexpression lines displayed a growth increase compared to GFP-overexpressing parasites, this was observed to be lower in glucosamine-treated PfHDAC1-GFP-glmS lines. Thus, even minor depletion of the overexpressed PfHDAC1 by glucosamine was found to be associated with loss of growth benefits.

We then tested if the overexpression of PfHDAC1 was associated with changes in progression of the cell cycle across the 48 hr. IDC by sampling parasites from the tightly synchronised PfHDAC1-GFP-glmS overexpression line and GFP-glmS control line at 8 hr intervals starting 10 hours post invasion (HPI). No significant changes in morphology or shift in phases of development were observed during PfHDAC1 overexpression vs the control line (Supp Fig. 3A). Thus, the increased parasite growth rate observed upon PfHDAC1 overexpression could not be explained by changes to parasite morphology or cell cycle progression. To gain insight into the transcriptional consequences of PfHDAC1 abundance in cells and to understand the phenotypic observations of the enhanced growth phenotype, we used strand-specific RNA sequencing to compare PfHDAC1-GFP-glmS overexpression vs GFP-glmS control overexpression parasites. RNA sequencing confirmed that PfHDAC1 was one of the top genes upregulated (log2 fold-change of 2.17; p-adjusted: 1.49e-55) in the episomal overexpression parasite lines. Interestingly, PfHDAC1 overexpression caused more genes to be upregulated (419 genes) than downregulated (35 genes) (log2fold-change >= -1/1; p-value<=0.05), indicating a strong indirect effect of PfHDAC1 overexpression on transcription (Fig. 4F). Among the downregulated genes, enrichment of biological processes was associated principally with Ca^2+^-mediated signaling and antigenic variation (∼9 genes). Moreover, Gene Ontology analysis of upregulated genes upon PfHDAC1 episomal overexpression revealed enrichment of genes for entry into host cells (rhoptry proteins, 6-cysteine proteins, merozoite surface protein), cytoadherence to microvasculature (EMP, rifin and stevor), immunomodulators (SERA 1, 3, 4 and 5), protein phosphorylation/signaling cascades (especially FIKK proteins) and exit from host cells (Fig. 4G and Supplementary Fig. 3B-E). Genes associated with host cell invasion/egress included merozoite surface protein (msp 1, 3 and 7), rhoptry neck protein (ron 3, 4 and 6) and high molecular weight rhoptry protein (rhoph2 and rhoph3), whose products help egressed parasites bind to fresh host RBCs during reinvasion. Thus, enhanced expression of these genes could be directly associated with better adhesion of egressed parasite forms (merozoites) onto fresh RBCs, enabling better propagation of infection over consecutive rounds of invasion.

### PfHDAC1 is downregulated in artemisinin-resistant lines, and its inhibition is associated with decreased sensitivity to artemisinin

Artemisinin-resistant parasites have been reported to modulate their transcriptional profile to reduce the activation of the drug and withstand the damage caused by it [43]. To compare the PfHDAC1 levels in artemisinin-resistant PfKelch13 C580Y mutant isolates and their artemisinin-sensitive WT counterpart parasites, we performed western blotting using anti-PfHDAC1 antibodies. We detected lower levels of PfHDAC1 protein in artemisinin-resistant parasites than in their sensitive counterparts (Fig. 5A). Thus, we corroborated that PfHDAC1 downregulation is indeed associated with artemisinin-resistant parasites. Interestingly, the skewed cell cycle dynamics (including persistent rings and smaller trophozoites) of romidepsin-treated parasites bear a strong resemblance to those of artemisinin-resistant parasites [44]. To minimize the activation and damage imposed by artemisinin, it is advantageous for parasites to lengthen the metabolically suppressed (relatively) ring stage and stunt trophozoite development [12]. Apart from the regulation of the cell cycle, several of the genes and associated biological pathways that we identified under PfHDAC1 regulation (heat shock proteins, DNA damage response, hemoglobin catabolism, and autophagy) are strongly implicated in artemisinin resistance [9, 11, 43, 45]. Since artemisinin-resistant parasites had lower levels of PfHDAC1, we were curious to know if the pharmacological inhibition of PfHDAC1 activity could tweak the artemisinin sensitivity of *P. falciparum*. To differentiate whether PfHDAC1 depletion in artemisinin-resistant lines was a downstream effect or a regulatory cause, we performed the ring-stage survival assay (RSA) to infer the sensitivity of parasites to artemisinin [46]. The baseline RSA of the tested lines was first established by calculating the percentage survival of parasites treated with dihydroartemisinin (DHA) relative to DMSO mock. Subsequently, to assess the effect of PfHDAC1 inhibition on artemisinin sensitivity, the parasites were exposed to DHA along with a sublethal dosage of an HDAC inhibitor (romidepsin/entinostat). The %RSA of the PfKelch13 wild-type *P. falciparum* 3D7 strain was found to be at an expected <1%. The artemisinin survival in parasites concomitantly treated with DHA and entinostat or romidepsin was reported at 4.5% and 5.6%, respectively, relative to the mock treatment (Fig. 5B). Thus, we found that PfHDAC1 inhibition during DHA exposure can enhance artemisinin resistance in otherwise drug-sensitive parasites. We further investigated the effect of PfHDAC1 inhibition on the drug sensitivity of various PfKelch13 mutant *P. falciparum* isolates. The C580Y mutant reported a baseline RSA of 4.2%, which increased marginally to 7.2% and 7% under DHA treatment concomitant with romidepsin or entinostat treatment, respectively (Fig. 5C). In another artemisinin-resistant *P. falciparum* Dd2 strain harboring a PfKelch13 R539T mutation, the RSA (baseline 9%) was increased to 11.4% under concomitant DHA and romidepsin treatment (Fig. 5D). We investigated whether the reduced parasite sensitivity to artemisinin could be explained (at least in part) by some of the transcriptional changes taking place upon PfHDAC1 inhibitor (romidepsin) treatment. Artemisinin-resistant parasites in the field exhibit upregulation of genes implicated in protein chaperoning, fatty acid biosynthesis, virulence, gametocytogenesis and phosphorylation/signaling. Pathways reported to be downregulated in resistant parasites, among others, include protein transport, endocytosis and RNA processing. An inspection of the genes deregulated upon romidespin treatment revealed the deregulation of these aforementioned pathways, thus providing some evidence that the PfHDAC1 inhibition-induced transcriptional response may mimic the artemisinin resistance transcriptome to some degree (Supplementary Fig. 4). Thus, our observations highlight that mild inhibition of PfHDAC1 activity can not only establish clinically significant artemisinin resistance in PfKelch13 wild-type isolates but can also enhance the level of resistance observed in already resistant PfKelch13 mutant strains.

**Figure 5:**
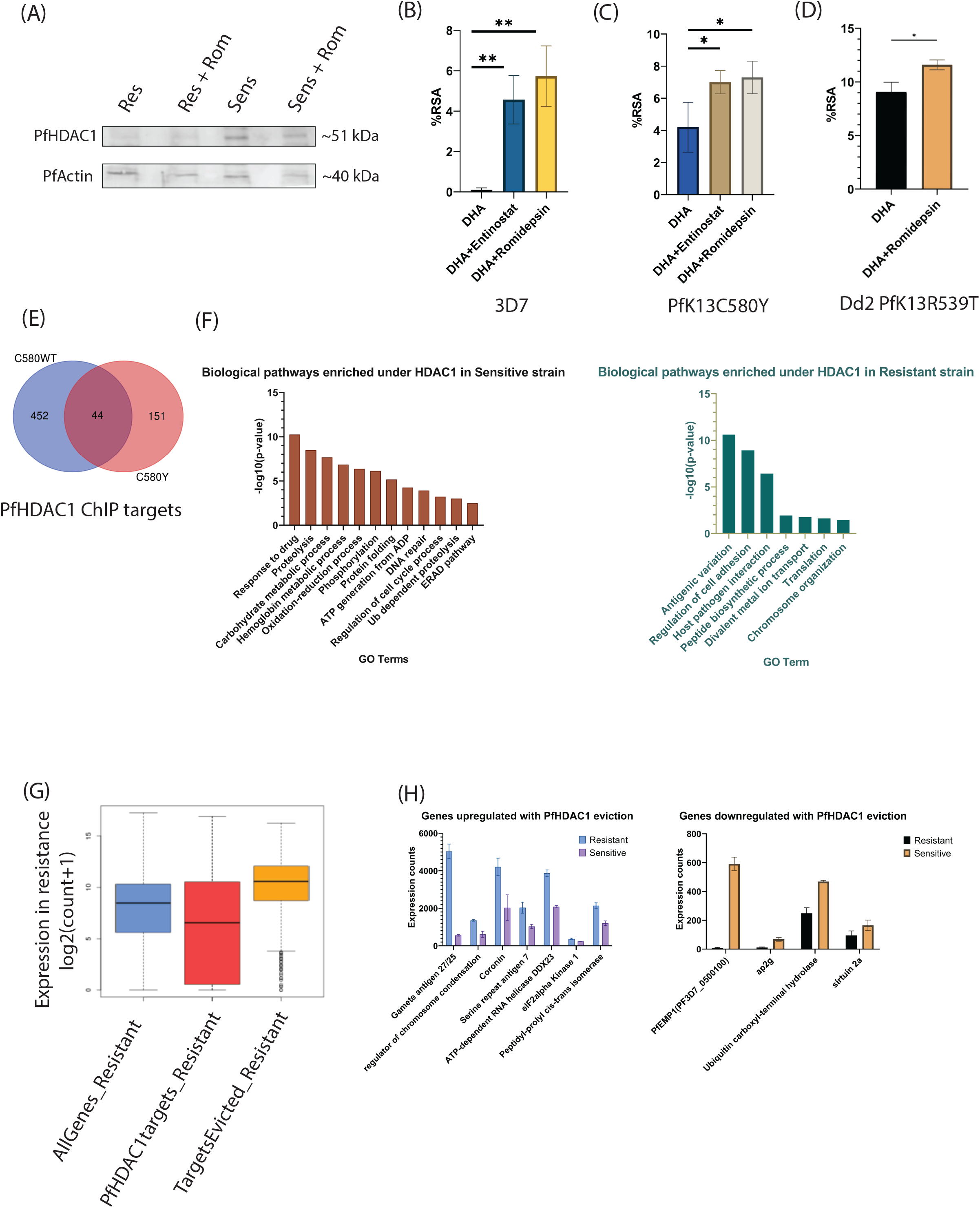
PfHDAC1 has lower abundance and differential genomic occupancy in artemisinin-resistant parasites. (A) Western blotting (α-PfHDAC1) confirmed that PfHDAC1 is depleted at the protein level in the artemisinin-resistant *P. falciparum* strain compared to its sensitive counterpart. Romidepsin treatment was administered to the respective parasite cultures to assess the effect of the inhibitor on protein levels. (B) Histograms representing the baseline %RSA of the artemisinin-sensitive *P. falciparum* 3D7 strain compared to the enhanced %RSA upon romidepsin and entinostat treatment. (Student’s unpaired t-test: Student’s unpaired t-test: Rom+DHA vs DHA alone, p-value=0.0032; Ent+DHA vs DHA alone, p-value=0.0032) (C) Histograms representing the baseline %RSA of artemisinin-resistant PfKelch13 C580Y strain compared to enhanced %RSA upon romidepsin and entinostat treatment. (Student’s unpaired t-test: Student’s unpaired t-test: Rom+DHA vs DHA alone, p-value=0.0435; Ent+DHA vs DHA alone, p-value=0.0396) (D) Histograms representing the baseline %RSA of artemisinin-resistant PfKelch13 R539T strain compared to enhanced %RSA upon romidepsin treatment. (Student’s unpaired t-test: Student’s unpaired t-test: Rom+DHA vs DHA alone, p-value=0.0213) (E) Venn diagram showing lower PfHDAC1 genomic targets in artemisinin-resistant vs sensitive strains. (F) Histograms for gene ontology enrichment analysis of biological pathways under PfHDAC1 regulation in (Left) artemisinin-sensitive and (Right) artemisinin-resistant parasites. (G) Boxplots representing genes with PfHDAC1 occupancy in resistant parasites show lower expression compared with pan transcriptome expression levels. However, PfHDAC1 gene targets in artemisinin-sensitive isolates that lose occupancy in resistant isolates report higher gene expression. (H) Histogram representing the expression of a subset of genes losing PfHDAC1 occupancy in PfKelch13C580Y mutant parasites. On the left are genes upregulated alongside loss of PfHDAC1 occupancy in resistant parasites; on the right are genes downregulated alongside loss of PfHDAC1 occupancy in resistant parasites.

Since artemisinin-resistant parasites show a reduction in PfHDAC1, we were interested in determining whether this could result in changes to and a possible reduction in its genomic occupancy between resistant and sensitive strains. We performed ChIP sequencing using an α-PfHDAC1 antibody in artemisinin-resistant (PfKelch13 C580Y) and artemisinin-sensitive (PfKelch13 C580WT) strains. We identified a total of 195 genes under PfHDAC1 regulation in the resistant strain compared to 496 genes under its regulation in the sensitive strain (Fig. 5E). Interestingly, 452 gene targets were lost from PfHDAC1 occupancy in resistant parasites, while 151 new targets were acquired. We believe that this reduced number of gene targets in resistant isolates is a consequence of lower PfHDAC1 abundance in the parasites. Gene Ontology analysis indicated that PfHDAC1 occupancy in sensitive strains was found on genes associated with response to drug, proteolysis, carbohydrate metabolism, hemoglobin catabolism, oxidation-reduction, phosphorylation, and cell cycle regulation (Fig. 5F, left). PfHDAC1 occupancy in resistant strains was mainly restricted to genes involved in antigenic variation and translation (Fig. 5F, Right). Considering that PfHDAC1 occupied a higher number of gene targets in the sensitive strain and lost a substantial fraction of those in resistant parasites, we wanted to see what happened to the expression of the genes from which PfHDAC1 lost occupancy in the resistant strain. We found that in the resistant parasites, PfHDAC1 target genes showed lower mean expression than the mean global gene expression. The genes that lost PfHDAC1 occupancy compared to the sensitive counterpart showed higher expression in artemisinin-resistant parasites (Fig. 5G). Thus, PfHDAC1 occupancy is associated with reduced gene expression on the genes in resistant strains, and its eviction from genes (originally bound in sensitive strain) elevates their expression. Among the genes that were significantly upregulated in the resistant strains after loss of PfHDAC1 occupancy were coronin, ATP-dependent RNA helicase DDX23, peptidyl-prolyl cis-trans isomerase, and eukaryotic translation initiation factor 2-alpha kinase (Fig. 5H). Among the genes downregulated with loss of PfHDAC1 occupancy were ubiquitin carboxyl-terminal hydrolase, a histone deacetylase sir2a, and erythrocyte membrane protein encoding var genes (PF3D7_1240300, PF3D7_1300100). Some of these genes are interesting since their deregulation or mutations have been reported to be associated with artemisinin resistance. Overall, we observed differential gene occupancy of PfHDAC1, with a lower abundance of the protein directly correlated with fewer genomic targets. The differential occupancy of PfHDAC1 in artemisinin-resistant vs artemisinin-sensitive strains may influence the expression of key genes associated with the emergence of artemisinin resistance.

### Artemisinin treatment inhibits phosphorylation and genomic occupancy of PfHDAC1

The antimalarial drug artemisinin is shown to target and form adducts with proteins belonging to several biological pathways in the parasite [47, 48]. We were curious to know if artemisinin treatment could affect the activity of PfCKII-α and its phosphorylation of PfHDAC1 in the kinase activity assay. We observed phosphorylation of PfHDAC1 in the absence of artemisinin (discussed in Fig. 1D). However, in the presence of 100 nM artemisinin, we observed a depletion in PfHDAC1 phosphorylation (Fig 6A; Lane 6). The phosphorylation of HDAC1 in mammalian systems has been shown to influence its catalytic activity as well as its interaction with other proteins and complex formation [37]. Protein-protein interactions and complex formation are crucial for the genomic targeting of transcription regulatory proteins such as HDACs. Our data already shows that depletion of phosphorylation on PfHDAC1 suppresses its catalytic activity, although we have not yet tested the effect of loss of PfHDAC1 phosphorylation on its interactome. We sought to investigate whether artemisinin exposure could also alter the genomic occupancy of PfHDAC1. We performed ChIP-seq for PfHDAC1 (using the PfHDAC1-2xFKBP-GFP knockin line) after 3 hours of 100 nM dihydroartemisinin treatment at approximately 21-24 HPI. This milder dosage and relatively shorter exposure time were chosen to avoid the consequences of rapid irrecoverable cell death. We found a significant reduction in the genomic occupancy of PfHDAC1 upon dihydroartemisinin exposure (464 genes) compared to the control condition (1409 genes) (Fig. 6B and 6C). A total of 988 target genes were lost, while 43 new target genes were acquired by PfHDAC1 upon dihydroartemisinin exposure. The pathways targeted by PfHDAC1 under control condition and lost under dihydroartemisinin exposure include splicing, hemoglobin catabolism, phosphorylation, redox homeostasis, cell cycle regulation, autophagy, vesicle fusion and protein ubiquitination. (Fig. 6D). On the other hand, PfHDAC1 targets under dihydroartemisinin treatment were associated with ornithine metabolism, purine containing nucleotide salvage and chromatin silencing pathways (Fig. 6D).

**Figure 6:**
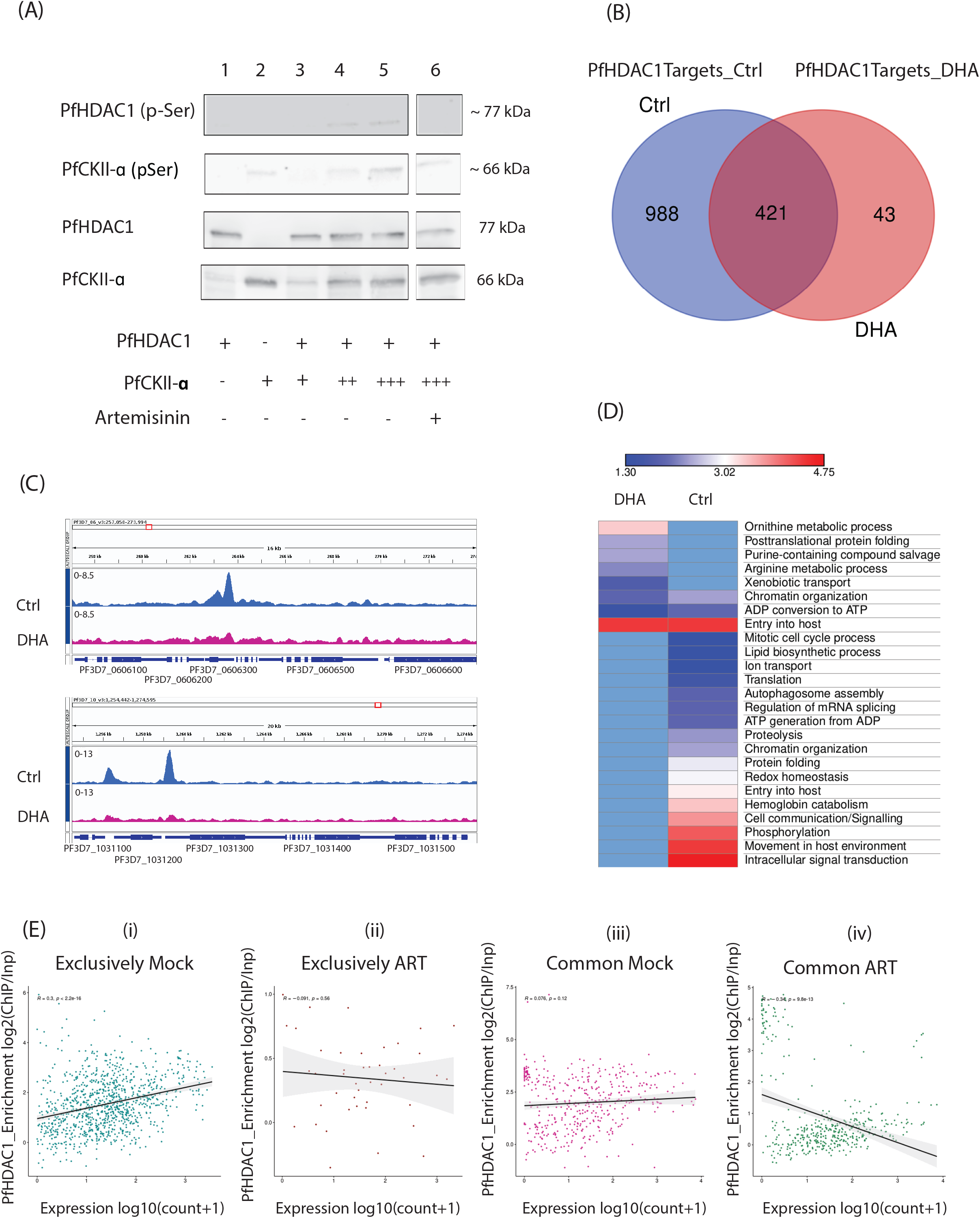
Artemisinin exposure interferes with PfHDAC1 genomic occupancy. (A) Western blot (using α-phospho Ser/Thr antibody in top two panels and α-GST antibody bottom two panels) for in vitro kinase activity assay using purified recombinant PfHDAC1 and PfCKII-α. The reaction mix components are as specified in the following table. Lanes 1-5 are the same as those in Fig. 1D; Lane 6 represents the reaction mixture spiked with artemisinin. (B) Venn diagram showing that dihydroartemisinin treatment interferes with the genomic occupancy of PfHDAC1, resulting in lower gene targets in the drug treatment than in the mock treatment. (C) Sample shots of PfHDAC1 ChIP-seq peaks in mock vs dihydroartemisinin treatment conditions (captured in IGV browser) highlighting the differential genomic enrichment of PfHDAC1 upon drug treatment. (D) Heatmap for gene ontology enrichment for distinct and shared biological processes under PfHDAC1 regulation in mock vs dihydroartemisinin treatment conditions. The scale bar on top is representative of -log10(p-value). (E) Scatter plots representing the correlation of PfHDAC1 occupancy (y-axis) vs gene expression on target genes. i) occupancy vs expression for genomic targets exclusive to mock condition (Pearson’s R=0.3; p-value=2.2e-16) ii) occupancy vs expression for genomic targets exclusive to artemisinin treatment condition (Pearson’s R= -0.091; p-value=0.56) iii) occupancy vs expression for common genomic targets in mock condition (Pearson’s R=0.076; p-value=0.12) iv) occupancy vs expression for common genomic targets in artemisinin treatment condition (Pearson’s R= -0.34; p-value=9.8e-13).

We further checked the correlation of PfHDAC1 genomic occupancy and gene expression from target genes in control and artemisinin treatment conditions. An in-house-generated RNA sequencing dataset was used for this purpose along with the PfHDAC1 ChIP sequencing datasets. We evaluated the expression of genes (in corresponding conditions) that were bound by PfHDAC1 exclusively under control or artemisinin treatment (Fig. 6 and Supplementary Fig. 5). For a subset of genes that were targets of PfHDAC1 regardless of treatment, we plotted the trend of binding vs expression in both control and artemisinin treatment. In general, under control/mock conditions, PfHDAC1 occupancy was found to be mildly positively correlated with the expression of bound genes (Pearson’s R=0.3; p-value=2.2e-16) (Fig 6E.i). These included genes belonging to pathways including splicing, lipid metabolism, haemoglobin catabolism, phosphorylation/signalling and redox homeostasis. On the other hand, for genes bound by PfHDAC1 exclusively under artemisinin treatment, we found no correlation between occupancy and expression (Pearson’s R= -0.091; p-value=0.56) (Fig 6E.ii). There was no significant enrichment for any biological pathways among these genes (except for a few genes encoding for putrescine metabolism). For the set of PfHDAC1 target genes in both mock and artemisinin-treated cells, occupancy was neutrally correlated with gene expression in the mock condition (Pearson’s R=0.076; p-value=0.12) and negatively correlated with expression under artemisinin treatment (Pearson’s R= -0.34; p-value=9.8e-13) (Fig 6E.iii and 6E.iv). These genes included those involved in host cell entry, protein folding and chromatin organization.

Overall, artemisinin treatment was found to suppress the genomic association of PfHDAC1 and dissociate the correlation of its genomic occupancy with target gene expression. Thus, artemisinin treatment possibly weakens the gene regulatory functions under PfHDAC1.

## Discussion

In this study, we focus on the class I histone deacetylase PfHDAC1, which is a prime antimalarial target [22, 23] and is critical for parasite viability [19, 23, 34]. We identified that the enzymatic activity of PfHDAC1 is governed by its posttranslational modification (serine phosphorylation) status, which is similar to what is observed for higher eukaryotic HDACs [37]. The phosphorylation PTM adds an important layer towards the regulation of its activity. We show that artemisinin treatment can inhibit the phosphorylation of PfHDAC1 and interfere with its genomic occupancy. Upon artemisinin exposure, PfHDAC1 evicts target sites associated with hemoglobin catabolism, phosphorylation/signaling, splicing, cell cycle regulation, autophagy, and proteostasis. Future experiments will be aimed at dissecting whether PfHDAC1 phosphorylation directly governs its genomic target selection. While these changes to genomic occupancy are induced upon drug treatment in sensitive parasites, over chronic exposure of parasites to the drug in the field, the changes to the genomic occupancy may become hard-wired into the parasites, leading to evolution of resistant forms. This is especially interesting in light of recent studies, which may indicate that the early transcriptional response to artemisinin in sensitive parasites may drive the mechanism of resistance emergence [49, 50].

Using class-specific HDAC inhibitors, we observed a deceleration of parasite progression through the IDC. This was also found to correlate with distortion in proper morphological development (especially DNA replication) and carry over of the infection to the next cycle. Using RNA sequencing, we identified the effects of romidepsin treatment on parasites, manifested in the deregulation of key genes associated with a wide array of biological processes, including RNA splicing, translation, transcriptional regulation, the cell cycle, DNA replication, and parasite interaction with host cells (entry/egress). Improper DNA replication caused by downregulation of associated genes may be responsible for insufficient conversion of “DNA light” forms of parasites (rings and early trophozoites) into “DNA heavy” forms resulting in faulty maturation. The episomal overexpression of PfHDAC1 was found to attribute growth advantage to parasites over consecutive invasion cycles. This was not surprising since HDAC1 overexpression has been shown to contribute to hyperproliferation phenotypes in mammalian systems [51]. RNA-sequencing of PfHDAC1-GFP-glmS episomal overexpression lines, however, did not reveal changes in genes related to the cell cycle or DNA replication but showed upregulation of genes associated with host cell invasion/egress, cytoadherence and several genes involved in the phosphorylation cascade. Through a higher abundance of invasion/egress-associated gene products, PfHDAC1 overexpression may account for this growth advantage [52]. Among the upregulated kinases, FIKK family members were observed in good numbers and have also been implicated in host cell remodelling and host-parasite interactions [53]. Future experiments should focus on testing host cell invasion and cytoadherence/rosetting by parasites upon PfHDAC1 overexpression. It is notable that the transcriptional changes observed with PfHDAC1 inhibitor romidepsin treatment and PfHDAC1 overexpression do not mirror each other dramatically. This may be due to off-target effects of the inhibitor treatment.

The effect of PfHDAC1 inhibition on the cell cycle mimics the cell cycle dynamics of artemisinin-resistant parasites, especially the slower IDC progression of a fraction of parasites. Artemisinin-resistant strains could have offset the effects of artemisinin by evolving for reduced levels of PfHDAC1. This would allow for slower growth and offer a protective advantage from the damage caused by artemisinin. We do note that the persistence of rings is only observed for a small fraction of parasites treated with inhibitors, which could be a reason behind the small increase in %RSA. The change to %RSA upon romidepsin treatment is still lower for PfKelch13 mutants (which have even lower PfHDAC1 to target). While PfHDAC1 pharmacological inhibitors are highly effective in inhibiting *Plasmodium* growth and are promising antimalarial compounds, we used sublethal dosages of the inhibitors. It is interesting to note how treatment with lower than lethal concentrations of these compounds can skew the parasite cell cycle enough and influence sensitivity to other drugs (artemisinin). We additionally analyzed the PfHDAC1 knockdown RNA-sequencing data generated by Huang et al. and identified that PfHDAC1 depletion is accompanied by broad-scale gene expression changes (Supp Fig. 2F) [34]. The expression of genes related to protein translation, chaperone, and fatty acid biosynthesis tended to increase, while genes related to the cell cycle, host invasion, phosphorylation, endocytosis and cytoadherence tended to be downregulated. Much of the transcriptional profile associated with targeted PfHDAC1 depletion matched the profile that is classically associated with resistance in Southeast Asia [11]. Thus, PfHDAC1 deregulation could drive gene expression patterns in parasites that allow for the emergence of resistance. It is worth noting that not all artemisinin-resistant parasites display downregulation of PfHDAC1. In such scenarios, alternative regulatory mechanisms may be in place. Thus, deregulation of PfHDAC1 may be a means to acquire resistance if not the only one to do so. As recognised by pioneering studies in *Plasmodium* drug resistance, processes that allow for a slower activation of artemisinin and compensate for the damage caused can modulate artemisinin sensitivity. PfHDAC1, with its regulation of several critical pathways, could be one such regulatory element that could fine tune the parasite’s responses to artemisinin, especially by skewing its cell cycle.

## Material and Methods

### Cloning, overexpression, and purification of recombinant PfHDAC1 and PfCKII-α

The full-length PfHDAC1 sequence was PCR amplified and cloned into the pET28a+ bacterial expression vector (NheI and XhoI restriction enzyme sites) and into the pGEX-4T1 expression vector (BamHI and XhoI sites) to obtain a 6xHis-tagged and GST-tagged version of the gene, respectively. The overexpression constructs were transformed independently into the BL21 (DE3) E. coli expression system. Bacterial culture growing at 0.5 optical density at 600 nm was induced with 0.5 mM isopropyl-1-thio-β-d-galactopyranoside IPTG for 5 hrs at 25 □. The bacterial pellet was resuspended in sonication buffer (10 mM Tris-Cl pH 8.0, 150 mM NaCl, 10% glycerol, 1XPIC and 1XPMSF) followed by sonication using a probe sonicator (Thomas Scientific, 70% amplitude, 10 minutes, 02 Sec on/06 Sec off). After sonication, the lysate was centrifuged at a high speed of 14,000 RPM for 30 minutes at 4°C. The supernatant was stored at -80°C, and the pellet was resuspended in 8 M urea prepared in sonication buffer. The pellet was kept under shaking conditions for 1 hour at room temperature followed by centrifugation. The earlier supernatant and pellet supernatant were run on a 10% resolving SDS PAGE gel along with uninduced bacterial lysate to check the induction of protein expression. Protein expression was also confirmed using western blotting (anti-His or anti-GST antibody). Once the expression was confirmed, protein purification was performed. GST-tagged PfHDAC1 was purified from the soluble fraction with glutathione Sepharose 4B beads (GE Healthcare) using 20 mM reduced glutathione. The 6xHis-tagged protein was purified from the insoluble fraction of a separately induced culture using Ni-NTA beads (GE Healthcare) with 250 mM imidazole. Purification of proteins was checked using SDS PAGE and western blotting. Purified proteins were dialyzed and stored at -20□.

PfCKII-α was PCR amplified from genomic DNA and cloned into the pEX4T1 vector for bacterial expression. The GST-tagged PfCKII-α clone was transformed into BL21 DE3 *E. coli* competent cells. Protein induction was performed with 1 M IPTG (at OD 0.6), and protein expression was allowed for 12 hrs. at 18°C. Cell lysate was prepared as described for PfHDAC1-GST tagged above, and purification of protein was performed using glutathione beads. The purified protein was eluted with 10 mM reduced glutathione and subsequently dialyzed.

### Antibodies

α-actin (Sigma A2066) and α-rabbit IgG (OSB PM035) were used for western blotting and immunoprecipitation, respectively. Goat α-rabbit Alexa Fluor 647 (Invitrogen A21245), goat α-rat Alexa Fluor 488 (Invitrogen A11006), and goat α-rabbit Alexa Fluor 488 (Invitrogen A11034) were used for immunofluorescence. Additionally, α-H3K9ac (Millipore 06-942), α-his (Biobharati Life Sciences BB-AB0010), α-GST (Cloud Clone Corp.) and α-phosphoserine/threonine (Abcam ab17464) were also used for western blotting. Rabbit polyclonal antibodies against PfHDAC1 were generated by immunizing rabbits with purified his-tagged recombinant PfHDAC1 (conjugated to Freund’s incomplete and complete adjuvant). New Zealand White rabbits (3-4 months old) were used for antibody generation. Two hundred micrograms of protein was intramuscularly injected for the first round, followed by 4 booster doses of 150 µg each. Five bleeds were collected via femoral bleeds and centrifuged to obtain the antisera from whole blood. The antibodies were affinity purified using GST-tagged recombinant PfHDAC1 conjugated to sulfolink resin (Thermo catalog 20402).

### *In vitro* protein-protein interaction assay

The recombinant purified PfHDAC1 and PfCKII-α were incubated together in an interaction buffer overnight for 4 hours at 4 □. The incubation reaction was split into two sets. Immunoprecipitation with anti PfHDAC1 antibodies was performed for one set and IgG for another at 4 [overnight (constant rolling). These served as the test and control pull down, respectively. The next day, washed recombinant protein-G Sepharose beads were added to the two reactions and incubated on rolling for 4 hours at 4 [. The bound complex was washed and eluted from the beads using 1 M glycine (pH 2.5) and neutralized with Tris pH 8.8. The complex was resolved on a 10% resolving SDS-PAGE gel, and western blotting was performed using an anti-GST antibody to identify the presence of tagged coimmunoprecipitated proteins.

### *In vitro* kinase activity assay

Highly purified recombinant PfHDAC1 and PfCKII-α were used as substrates and enzymes for the *in vitro* kinase assay. PfHDAC1 (500 ng) was incubated with increasing amounts (360 ng, 780 ng, 1080 ng) of PfCKII-α in the presence of a kinase activity buffer (20 mM Tris HCl pH 7.5, 20 mM MgCl2, 2 mM MnCl2, 0.1 mM PMSF) with 10 µM ATP. For a baseline control, PfHDAC1 incubated without PfCKII-α was taken. Furthermore, two reactions with increasing dosages of the PfCKII-α-specific inhibitors 4,5,6,7-tetramethylbromobenzotriazole and 4,5,6,7-tetramethylbromo-2-azabenzimidazole (TBB) (Merck/Calbiochem CAS 17374-26-4) were performed to check for possible suppression of phosphoryl group transfer. Furthermore, to probe the effect of artemisinin treatment on the kinase reaction, we set up an additional reaction with 100 nM artemisinin. The reaction was incubated at 37 [on a thermal mixer and then run on a 10% resolving SDS-PAGE gel followed by western blotting with anti-phospho-serine antibody.

### *In vitro* deacetylase activity assay

Highly purified recombinant PfHDAC1 was first phosphorylated in the *in vitro* kinase assay. Phosphorylated PfHDAC1 was used as the enzyme, and histones isolated from the *P. falciparum* pellet were used as substrates for this reaction. Two micrograms of histones was incubated with increasing amounts of PfHDAC1 in the presence of deacetylase buffer (25 mM Tris HCl, pH 8.0, 137 mM NaCl, 2.7 mM KCl, 1 mM MgCl2, 0.1 mM ZnCl2). For the control reaction, only purified histones were incubated in the deacetylase reaction mix (without PfHDAC1). The class I HDAC-specific inhibitor entinostat was used in an additional reaction to inhibit the potential deacetylase activity of PfHDAC1. PfHDAC1 obtained from the kinase assay (incubated with kinase inhibitor) was used as a nonphosphorylated enzyme and essentially served as a control. After incubation at 37 □ for 1 hr. The reactions were run on a 15% resolving SDS-PAGE gel, and western blotting was performed using anti-H3K9ac antibody.

### Generation of GFP knockin constructs for PfHDAC1

For the GFP knockin construct, 750 bp of the C-terminal region of PfHDAC1 was cloned into the parent pSLI-2xFKBP-GFP plasmid generated by the Tobias Spielmann team (deposited to Addgene).

### *P. falciparum* transfections

Lonza nucleofector 4D was used for transfection of plasmids into highly synchronized parasites. Eighty micrograms of knockin plasmid was dissolved in 100 µl of P3 primary solution. Double synchronised (Percoll centrifugation) segmented schizonts were mixed with the DNA/P3 solution mix and nucleoporated in the nucleofector 4D machine with the pulse program FP 158 (designed for *P. berghei*). The zapped cells were immediately transferred to a T25 flask with 3 ml of media and 200 µl fresh RBCs and placed on a shaker incubator at 37°C for 2 hours to allow the egressed merozoites to invade the fresh cells. The flask was later supplemented with additional media to make it 2% hematocrit. Drug selection was started 24 hours post transfection. The appearance of transgenic lines was checked initially via Giemsa smear.

### Cloning of the transgenic parasite lines

The validated transgenic lines were serially diluted in a 96-well flat plate to obtain 200 µl cultures with 1 parasite per well. These were confirmed with plaque formations and then transferred onto round bottom 96-well plates for expansion. The cells were then microscopically confirmed for GFP expression and expanded into flasks.

### alidation of the PfHDAC1-2xFKBP-GFP transgenic line

Confocal microscopy and western blotting using anti-GFP primary antibody were used to validate the PfHDAC1-2xFKBP-GFP transgenic line. The transgenics were generated and tested at Dr. Moritz Treeck’s laboratory at The Francis Crick Institute, London.

### Chromatin immunoprecipitation

The PfHDAC1-2xFKBP-GFP knockin NF54 line parasitized RBCs (mock or 100 nM dihydro artemisinin treated from ∼21-24 HPI/3 hours) were crosslinked using 1% formaldehyde (Thermo Scientific, 28908) for 10 mins at RT. ChIP was also performed in artemisinin-resistant MRA-1236 and artemisinin-sensitive MRA-1254 (crosslinked at approximately 24HPI). Then, 150 mM glycine was added to quench the cross-linking reaction. The samples were washed using 1X PBS (chilled) before proceeding with lysis. Sample homogenization was performed using swelling buffer (25 mM Tris-Cl pH 7.9, 1.5 mM MgCl2, 10 mM KCl, 0.1% NP-40, 1 mM DTT, 0.5 mM PMSF, 1xPIC) followed by cell lysis in sonication buffer (10 mM Tris-Cl pH 7.5, 200 mM NaCl, 1% SDS, 4% NP-40, 1 mM PMSF, 1X PIC). Sonication was performed using Covaris S220 to obtain chromatin sizes of 200-400 bp. Preclearing was performed for 1 hr at 4°C using recombinant protein G-conjugated Sepharose beads with continuous gentle inverting. Thirty micrograms of purified chromatin was used per immunoprecipitation reaction (α-GFP antibody for the knockin line and α-PfHDAC1 antibody for the resistant and sensitive lines) and incubated for 12 h at 4 [. Samples were then incubated with saturated Protein G Sepharose beads for 4 hours at 4 [. Bound chromatin was finally washed with low salt, high salt, LiCl wash buffers followed by TE buffer wash and eluted using ChIP elution buffer (1% SDS, 0.1 M sodium bicarbonate). Both the ChIP and input samples were reverse crosslinked using 0.3 M NaCl overnight at 65 [along with RNase A treatment. Proteinase K treatment was performed at 42 [the next day for 1 hour. Finally, DNA was purified using phenol chloroform isoamyl alcohol precipitation.

### ChIP-sequencing library preparation and sequencing

ChIP-sequencing libraries were prepared from 5-10 ng of DNA using the NEBNext Ultra II DNA Library Prep kit. Chromatin was immunoprecipitated, fragmented DNA samples were end repaired, and adapters were ligated. Size selection was performed using Agencourt XP beads (Beckman Coulter). Adapter-ligated fragments were PCR amplified using indexing primers followed by purification using Agencourt XP beads (Beckman Coulter). The library electropherograms were assessed using an Agilent Bioanalyzer 2100 and Agilent DNA 1000 kit. The sequencing for these was done at the Next Generation Sequencing facility at IISER, Pune using the Illumina NextSeq 550 sequencer using 2×150 bp specs for the PfHDAC1-2xFKBP-GFP knockin NF54 line (mock and ART) and 75 bp for MRA-1236 and MRA-1254 lines.

### ChIP-Seq data analysis

The sequencing reads were demultiplexed using the bcl2fastq tool in the Linux platform. Read quality checks were performed using FASTQC, and sub-quality reads (not passing the Q30 filter) and adaptor sequences were trimmed using Trim_Glaore. Reads were aligned onto the *Plasmodium falciparum* 3D7 reference genome using Bowtie aligner. SAMTools was used to further process the data (sorting, etc.). The sorted ChIP and input BAM files were then used for peak calling in MACS2 software. Background subtraction was performed using the bdgcmp command in MACS. Peak annotation was performed using Bedtools, and the peaks were visualized using the IGV genome browser [54, 55]. The SeMonk tool was used to compare the tag densities across the two ChIP replicate experiments on the PfHDAC1-2xFKBP-GFP knockin line to validate the correlation between the two.

The DeepTools suite was used for additional ChIP data analysis and visualization [56]. The input normalized fold enrichment bedgraph files were converted into bigwig format. The computeMatrix option was used to generate an enrichment matrix for PfHDAC1 over the average gene body flanked by 500 bp of upstream and downstream regions. The matrix was then fed to the plotHeatmap tool to generate the PfHDAC1 occupancy heatmap on the target genes. The PlosDB Gene Ontology tool was used to identify the biological pathways enriched for PfHDAC1 occupancy gene targets [57]. To correlate PfHDAC1 occupancy with gene expression in mock and ART treatment, ggplot2 was used to plot the ChIP occupancy vs gene expression counts from the relevant RNA sequencing dataset. We plotted for gene targets exclusively bound by PfHDAC1 in control/mock and artemisinin treatment and a subset of genes bound by PfHDAC1 in both conditions with the relevant RNA sequencing data.

### PfHDAC1-specific inhibitor treatment for RT-qPCR and protein analysis

Parasite cultures grown at 5% parasitemia and 2% hematocrit were treated _with a 2x IC50_ dosage of entinostat or romidepsin for 4 hours prior to harvesting for gene expression analysis using RT-qPCR. The same treatment was carried out for 8 hours prior to harvesting to estimate the effect on protein levels with western blotting.

### Quantitative RT-PCR

Two micrograms of DNAse-free RNA was used for cDNA synthesis using an InProm-II Reverse transcription system (Promega) according to the manufacturer’s recommendation. Random primers were used for cDNA synthesis. Real-time PCR was carried out using a CFX96 Real Time PCR detection system (Bio-Rad). 18S rRNA and seryl-tRNA synthetase were used as internal controls to normalize for variability across samples. Quantification of the expression was performed with the help of a fluorescence readout of SYBR green dye incorporation into the amplifying targets using a Bio-Rad thermocycler. Each experiment included three technical replicates and was performed for three independent biological replicates.

### Investigation of the effect of PfHDAC1 inhibition on the intraerythrocytic developmental cycle

Highly synchronized (Percoll density gradient centrifugation) parasites were cultured at ∼5% parasitemia and 2% hematocrit and treated with 0.5xIC_50_ concentration of PfHDAC1 inhibitor romidepsin) at approximately 6 hours post invasion (regarded as 0 hr post treatment). The parasites were then sampled every 8 hours for the next 48 hours and frozen for either SYBR fluorometry to estimate the DNA content or prepared into thin Giemsa-stained smears for microscopic examination of morphology and stage progression. The SYBR fluorometry samples were lysed with SYBR Green I lysis buffer and processed for DNA signal intensity via fluorometry (excitation: 495 nm; emission: 520 nm). The relative DNA signal for inhibitor treatment estimated by fluorescence was plotted as a percentage against mock sample readouts. For parasite proliferation estimation, parasitemia was recounted upon completion of one cycle in the mock-vs inhibitor-treated samples. The relative percentage of parasites from each of the developmental stages of *P. falciparum* was calculated and plotted on histograms. Each time point was validated with at least 1000 cell count observations.

### Generation of overexpression constructs for PfHDAC1

The full-length sequence of PfHDAC1 was PCR amplified from genomic DNA using sequence-specific primers and cloned into the pDC2 overexpression vector using the AvrII and NheI restriction sites. This put PfHDAC1 under the control of the calmodulin promoter and in frame with a C-terminal GFP tag. Furthermore, to add a layer of regulatability to the overexpression system, we cloned a glmS ribozyme sequence (PCR amplified from pHSP101 plasmid) in frame with the GFP tag using the XhoI enzyme. This resulted in an overexpression system synthesizing the PfHDAC1-GFP-glmS fusion transcript.

### *P. falciparum* transfections for PfHDAC1-GFP-glmS overexpression constructs

Transfections for the control (GFP-glmS) and test (PfHDAC1-GFP-glmS) were performed in the artemisinin-sensitive PfKelch13WT (MRA-1252 parasite line sourced from the MR4 repository). The transfection protocol was followed as described previously except that 30 µg of overexpression plasmid was used. Drug selection was started 24 hours posttransfection with 2 µg/ml blasticidin-S. Confirmation of GFP expression and clonal selection of transgenic parasites was performed as described previously for the PfHDAC1-2xFKBP-GFP knockin line. The transgenics were generated and tested at Dr. Moritz Treeck’s laboratory at The Francis Crick Institute, London.

### Validation of reversible overexpression of PfHDAC1-GFP

Validation of PfHDAC1-GFP overexpression and reversible depletion was validated with treatment of culture with 0 and 2.5 mM glucosamine-HCl followed by western blotting for depletion of GFP-tagged PfHDAC1 signal.

### Parasite growth curve assay

GFP-glmS ctrl overexpression and PfHDAC1-GFP-glmS overexpression were tightly synchronized using two rounds of Percoll density gradient centrifugation. Parasitemia was estimated by staining cells with SYBR Green I dye and subjecting them to flow cytometry. The cultures were diluted to 1% starting parasitemia and 2% hematocrit in 96-well plates and allowed to proliferate over a duration of 6 days/3 IDC. To test the effect of glucosamine dosage (and episomal expression suppression) on the growth trend, wells with 2.5 mM glucosamine treatment were set up in parallel with mock-treated parasites for both PfHDAC1-GFP-glmS overexpression and GFP-glmS overexpression lines. Media was replenished carefully after the third day with accurate replacement for mock and drugs. Each strain and condition were set in triplicate on a 96-well plate. The culture in the wells was sampled every 24 hours until the end of the assay and subjected to SYBR Green I flow cytometry to calculate parasitemia progression over the course of the experiment. Growth curves were plotted on GraphPad Prism software. The baseline toxicity of glucosamine was calculated based on the viability of the GFP-glmS lines and then subtracted from the cognate readouts of PfHDAC1-GFP-glmS to represent toxicity-corrected GFP-subtracted growth curves.

### Ring-stage survival assay

Tightly synchronised (double Percoll density gradient centrifugation) 0-3 hr 3D7 *P. falciparum* ring stage parasites were taken at 1.5% parasitemia and 2% hematocrit in 200 µl volume 96-well plates. Parasites were then either treated with DMSO mock or with 700 nM dihydroartemisinin for a duration of 6 hrs, after which the mock/drug was washed off and the parasites were reinstated in culture. They were allowed to grow for another 66 hrs. At the end of 72 hours, live parasites were counted using Giemsa smears for at least 10,000 cells in both mock and DHA-treated cultures. The percentage of cells surviving DHA treatment relative to the mock treatment was logged as % RSA. To check the effect of PfHDAC1 inhibition, tightly synchronised freshly invaded ring stages were treated with 0.25x IC_50_ dosage of the selected PfHDAC1 inhibitor and then half an hour later followed by DHA or mock treatment, which was continued for 6 hrs. The drugs/mock were washed off at the end of 6 h of DHA treatment; however, the inhibitors were readded to the culture at the original conc. until the completion of the assay at 66 hr post DHA wash off. The percentage of cells surviving concomitant inhibitor + DHA treatment relative to those surviving inhibitor + mock treatment was taken as the %RSA. The assays were carried out in technical triplicates and repeated for two biological replicates.

### Culture for RNA sequencing of PfHDAC1 overexpression and PfHDAC1 pharmacological inhibition experiments

For strand-specific RNA sequencing, parasites carrying the control GFP-glmS episomal construct and PfHDAC1-GFP-glmS episomal construct were cultured in triplicate at 8% parasitemia and 2% hematocrit. After double synchronization, the cultures were harvested at approximately 24 hours post invasion. For strand-specific RNA sequencing upon mock vs romidepsin treatment, NF54 strain parasites were cultured in triplicate at 8% parasitemia and 2% hematocrit. After double synchronization, one set of replicates was treated with DMSO mock, while the test set of replicates was treated with 360 nM romidepsin (2XIC_50_) for 3 hours at 21 HPI and harvested at 24 hours post invasion.

### RNA isolation and RNA sequencing library preparation

After harvesting the parasites, we proceeded with the isolation of RNA from the TriZol suspended samples. The Thermo Colibri 3’ mRNA library preparation kit was utilized for the preparation of strand-specific libraries. The final amplified libraries were quality checked on an Agilent bioanalyzer for size distribution and Qubit fluorometer for quantity. Single-end sequencing was performed on the Illumina NextSeq 550 platform (150 bp read length).

### RNA sequencing and data analysis

The sequencing data were quality checked using FASTQC and trimmed for low quality (Q30) and adaptor sequences. The trimmed reads were then mapped onto the *Plasmodium falciparum* 3D7 genome ver. 53 using Hisat2 (unpaired reads and reverse stranded orientation settings). HTSeq-count was used to quantify the raw reads from the mapped datasets, and then DESeq2 was used for sample-wise read normalization and differential gene expression analysis with the cognate test vs ctrl samples (PfHDAC1 overexpression vs GFP overexpression; romidepsin vs mock treatment). Figures and plots were generated with the R environment and GraphPad Prism software.

### Additional experiments and RNA sequencing datasets used in the analysis

Double synchronized cultures of the PfKelch13C580Y and PfKelch13C580WT isolates (MRA1236/Cam2 and MRA1254/Cam2_rev, respectively) were maintained at 5% parasitemia and 2% hematocrit. Harvesting for RNA was performed at 24 HPI. RNA isolation was performed using a standard TRIzol reagent protocol. One microgram of total RNA was used for library preparation. mRNA was enriched from total RNA using the Agilent SureSelect mRNA stranded library preparation kit. The stranded RNA sequencing library was subsequently prepared per the manufacturer’s protocol. The library was sequenced in 2×150 bp paired-end format using the Illumina NextSeq550 platform.

For mock vs art treatment experiments, tightly synchronized *Plasmodium falciparum* 3D7 parasites were cultured at 5% parasitemia and 2% hematocrit. Parasites were then treated with either DMSO mock or 30 nM artemisinin for 6 hours at the late ring stage and harvested at 23HPI. RNA isolation was performed using a standard TRIzol reagent protocol. One microgram of total RNA was used for library preparation. The Agilent SureSelect mRNA stranded library preparation kit protocol was followed. The library was sequenced in 2×150 bp paired-end format using the Illumina NextSeq550 platform.

The sequencing dataset was trimmed for low-quality bases (q30 cut-off) and adaptor sequences using Trim_Galore. The fastq files were then aligned onto *Plasmodium falciparum* genome ver. 53 for mock vs DHA treatment datasets and ver. 41 for the MRA1236 and MRA1254 isolates using the Hisat2 aligner. HTSeq-count was used to generate count data for the gene features.

### Data access

ChIP-sequencing data for PfHDAC1 as well as gene expression data (RNA sequencing) for different conditions are submitted to the Sequence Read Archive (SRA) under BioProject ID PRJNA817874.

## Ethics Statement

This study does not involve human participants. The human RBCs used in this study were obtained from the KEM Blood Bank (Pune, India) as blood from anonymized donors. The use of rabbits in this study for immunization (IISER/IAEC/2017-01/008) was reviewed and approved by IISER-Pune Animal House Facility (IISER: Reg No. 1496/GO/ReBi/S/11/CPCSEA). The approval is as per the guidelines issued by the Committee for the Purpose of Control and Supervision of Experiments on Animals (CPCSEA), Govt. of India.

## Acknowledgements

The following reagents were obtained through BEI Resources (www.mr4.org), NIAID, NIH: *Plasmodium falciparum*, Strain IPC 3445 (MRA-1236) contributed by Didier Menard and *Plasmodium falciparum*, Strain Cam2_rev (MRA-1254) contributed by David A. Fidock. The parasite line (MRA-1252) used for PfHDAC1 overexpression was obtained from the London School of Tropical Health and Medicine originally sourced from MR4 (contributed by Didier Menard and David Fidock). This work was supported by the DBT-Genome Engineering Technologies program (BT/PR25858/GET/119/169/2017) from the Government of India to KK. AK is supported by the CSIR-SPM fellowship. AK is also supported by the EMBO Scientific Exchange Grant (#8640) to carry out part of this work. MT and HD received funding from the Francis Crick Institute which receives its core funding from Cancer Research UK (FC001189), the UK Medical Research Council (FC001189), and the Wellcome Trust (FC001189). The funders had no role in the study design, data collection and analysis, decision to publish, or preparation of the manuscript. We also would like to acknowledge the use of BioRender (online tool) images for the creation of Fig. 2(b).

## Authors’ contributions

AK designed and performed the experiments and analysed the data. HD contributed to the generation of the PfHDAC1-2xFKBP-GFP knockin and PfHDAC1-GFP-glmS overexpression lines and the design and performance of growth assay experiments. BD designed and performed experiments and analyzed data related to parasite growth and drug sensitivity assays. DF designed and performed the experiments and analyzed the data related to the PfCKII-α and PfHDAC1 interaction/kinase assays and the PfHDAC1 histone deacetylase activity assay. MT coordinated and supervised the generation of transgenic parasite lines and experiments on them at his lab at the Francis Crick Institute, London. AK and KK wrote the manuscript. KK planned, coordinated, and supervised the project. All authors read and approved the final manuscript.

## Supplementary Figure Legends

**Supplementary Figure 1:** (A) Netphos 3.1 analysis of the PfHDAC1 amino acid sequence predicts CKII to be the kinase associated with S391, S397 and S440 phosphorylated residues. (B) FASTA format sequence of PfHDAC1 protein followed by the predicted posttranslational modifications at key sites. (C) Histogram representing the phosphorylation potential of serine, threonine and tyrosine residues across the PfHDAC1 sequence. Serine residues 391 and 397 are boxed in black, while serine 440 is boxed in blue for clarity.

**Supplementary Figure 2:** (A) SDS-PAGE and (B) α-GST western blotting for purified recombinant PfCKII-α-GST. (C) Western blotting (α-GST) for the in vitro interaction assay between purified recombinant PfCKII-α and PfHDAC1. (D) Western blot (α-H3K9ac) for the in vitro histone deacetylase activity assay using purified recombinant PfHDAC1 (phosphorylated or non-phosphorylated) as the enzyme and histones isolated from *P. falciparum* as substrates. The reaction mix components are mentioned in the following table. (E) Scatter plot comparing the tag counts of the two PfHDAC1 ChIP seq replicates showing a high degree of correlation between the two (R=0.945) (F) Boxplots representing genes belonging to artemisinin resistance-relevant biological pathways (Left) upregulated and (Right) downregulated with glucosamine-mediated depletion of PfHDAC1-GFP-glmS. Data analyzed from Huang et al., 2020; Cell Discovery.

**Supplementary Figure 3:** (A) Giemsa-stained smear panel comparing parasite progression through the intraerythrocytic development cycle for GFP ctrl overexpression and PfHDAC1 overexpression parasite lines. Histograms representing the gene upregulated with PfHDAC1 overexpression in parasites related to invasion (B), phosphorylation (C), and cytoadherence (D). Genes associated with mitochondrial ETC and Ca2+ signaling were found to be downregulated in the same dataset (E). The fold-changes were calculated by DESeq2 differential gene expression analysis and are supported by p-value <=0.05.

**Supplementary Figure 4:** Boxplots for genes belonging to specific biological pathways (A) upregulated and (B) downregulated with romidepsin treatment. Some of these pathways are reported to be deregulated in artemisinin-resistant parasites (highlighted in red). Wilcox statistical analysis was performed for the data in R (using ggplot2), indicating the p-values.

**Supplementary Figure 5:** Scatter plots representing the correlation of PfHDAC1 occupancy with target gene expression in (Left) mock-and (Right) artemisinin-treated parasites.

**Supplementary Table Document 1:** A table summarizing the ChIP-sequencing and RNA-sequencing experiments performed in this study with details on sample ID, strain and stage of parasite, treatment and duration (if any).

**Supplementary Table Document 2:** Summary of PfHDAC1 ChIP targets observed with >= 2-fold enrichment in mock vs dihydroartemisinin treatment conditions (Tabs 1 and 2, respectively). Tabs 3 and 4 list PfHDAC1 ChIP targets with >=2-fold enrichment in PfKelch13 C580WT (artemisinin sensitive) and PfKelch13 C580Y (artemisinin resistant) parasites, respectively.

**Supplementary Table Document 3:** Unfiltered summary of differential gene expression analysis performed using DESeq2 on PfHDAC1-GFP-glmS vs GFP-glmS parasites (Tab 1) and mock vs romidepsin-treated parasites (Tab 2).

## Bibliography

1. WHO (2020) World malaria report 2020: 20 years of global progress and challenges in World malaria report 2020: 20 years of global progress and challenges.

2. Ross, A., Maire, N., Molineaux, L. & Smith, T. (2006) An epidemiologic model of severe morbidity and mortality caused by Plasmodium falciparum, The American Journal of Tropical Medicine Hygiene. 75, 63–73.

3. Müller, I. B. & Hyde, J. E. (2010) Antimalarial drugs: modes of action and mechanisms of parasite resistance, Future Microbiology. 5, 1857–1873.

4. Menard, D. & Dondorp, A. (2017) Antimalarial drug resistance: a threat to malaria elimination, Cold Spring Harbor Perspectives in Medicine. 7, a025619.

5. WHO (2016) Artemisinin and artemisinin-based combination therapy resistance: status report in, World Health Organization,

6. Tilley, L., Straimer, J., Gnädig, N. F., Ralph, S. A. & Fidock, D. A. (2016) Artemisinin action and resistance in Plasmodium falciparum, Trends in Parasitology. 32, 682–696.

7. Ariey, F., Witkowski, B., Amaratunga, C., Beghain, J., Langlois, A.-C., Khim, N., Kim, S., Duru, V., Bouchier, C. & Ma, L. (2014) A molecular marker of artemisinin-resistant Plasmodium falciparum malaria, Nature. 505, 50–55.

8. Demas, A. R., Sharma, A. I., Wong, W., Early, A. M., Redmond, S., Bopp, S., Neafsey, D. E., Volkman, S. K., Hartl, D. L. & Wirth, D. F. (2018) Mutations in Plasmodium falciparum actin-binding protein coronin confer reduced artemisinin susceptibility, Proceedings of the National Academy of Sciences. 115, 12799–12804.

9. Birnbaum, J., Scharf, S., Schmidt, S., Jonscher, E., Hoeijmakers, W. A. M., Flemming, S., Toenhake, C. G., Schmitt, M., Sabitzki, R. & Bergmann, B. (2020) A Kelch13-defined endocytosis pathway mediates artemisinin resistance in malaria parasites, Science. 367, 51–59.

10. Mbengue, A., Bhattacharjee, S., Pandharkar, T., Liu, H., Estiu, G., Stahelin, R. V., Rizk, S. S., Njimoh, D. L., Ryan, Y. & Chotivanich, K. (2015) A molecular mechanism of artemisinin resistance in Plasmodium falciparum malaria, Nature. 520, 683–687.

11. Mok, S., Ashley, E. A., Ferreira, P. E., Zhu, L., Lin, Z., Yeo, T., Chotivanich, K., Imwong, M., Pukrittayakamee, S. & Dhorda, M. (2015) Population transcriptomics of human malaria parasites reveals the mechanism of artemisinin resistance, Science. 347, 431–435.

12. Mok, S., Stokes, B. H., Gnädig, N. F., Ross, L. S., Yeo, T., Amaratunga, C., Allman, E., Solyakov, L., Bottrill, A. R. & Tripathi, J. (2021) Artemisinin-resistant K13 mutations rewire Plasmodium falciparum’s intra-erythrocytic metabolic program to enhance survival, Nature Communications. 12, 1–15.

13. Toenhake, C. G. & Bártfai, R. (2019) What functional genomics has taught us about transcriptional regulation in malaria parasites, Briefings in Functional Genomics. 18, 290–301.

14. Bozdech, Z., Llinás, M., Pulliam, B. L., Wong, E. D., Zhu, J., DeRisi, J. L. & Ward, G. (2003) The transcriptome of the intraerythrocytic developmental cycle of Plasmodium falciparum, PLoS Biology. 1, e5.

15. Le Roch, K. G., Johnson, J. R., Florens, L., Zhou, Y., Santrosyan, A., Grainger, M., Yan, S. F., Williamson, K. C., Holder, A. A. & Carucci, D. J. (2004) Global analysis of transcript and protein levels across the Plasmodium falciparum life cycle, Genome Research. 14, 2308–2318.

16. Karmodiya, K., Pradhan, S. J., Joshi, B., Jangid, R., Reddy, P. C. & Galande, S. (2015) A comprehensive epigenome map of Plasmodium falciparum reveals unique mechanisms of transcriptional regulation and identifies H3K36me2 as a global mark of gene suppression, Epigenetics and Chromatin. 8, 1–18.

17. Kanyal, A., Rawat, M., Gurung, P., Choubey, D., Anamika, K. & Karmodiya, K. (2018) Genome[wide survey and phylogenetic analysis of histone acetyltransferases and histone deacetylases of Plasmodium falciparum, The FEBS Journal. 285, 1767–1782.

18. You, D., Richardson, J. R. & Aleksunes, L. M. (2020) Epigenetic regulation of multidrug resistance protein 1 and breast cancer resistance protein transporters by histone deacetylase inhibition, Drug Metabolism Disposition. 48, 459–480.

19. Engel, J. A., Jones, A. J., Avery, V. M., Sumanadasa, S. D., Ng, S. S., Fairlie, D. P., Adams, T. S. & Andrews, K. T. (2015) Profiling the anti-protozoal activity of anti-cancer HDAC inhibitors against Plasmodium and Trypanosoma parasites, International Journal for Parasitology: Drugs Drug Resistance. 5, 117–126.

20. Bushell, E., Gomes, A. R., Sanderson, T., Anar, B., Girling, G., Herd, C., Metcalf, T., Modrzynska, K., Schwach, F. & Martin, R. E. (2017) Functional profiling of a Plasmodium genome reveals an abundance of essential genes, Cell. 170, 260-272. e8.

21. Zhang, M., Wang, C., Otto, T. D., Oberstaller, J., Liao, X., Adapa, S. R., Udenze, K., Bronner, I. F., Casandra, D. & Mayho, M. (2018) Uncovering the essential genes of the human malaria parasite Plasmodium falciparum by saturation mutagenesis, Science. 360.

22. Elbadawi, M. A. A., Awadalla, M. K. A., Hamid, M. M. A., Mohamed, M. A. & Awad, T. A. (2015) Valproic acid as a potential inhibitor of Plasmodium falciparum histone deacetylase 1 (PfHDAC1): an in silico approach, International Journal of Molecular Sciences. 16, 3915–3931.

23. Hansen, F. K., Sumanadasa, S. D., Stenzel, K., Duffy, S., Meister, S., Marek, L., Schmetter, R., Kuna, K., Hamacher, A. & Mordmüller, B. (2014) Discovery of HDAC inhibitors with potent activity against multiple malaria parasite life cycle stages, European Journal of Medicinal Chemistry. 82, 204–213.

24. Hesping, E., Skinner-Adams, T. S., Fisher, G. M., Kurz, T. & Andrews, K. T. (2020) An ELISA method to assess HDAC inhibitor-induced alterations to P. falciparum histone lysine acetylation, International Journal for Parasitology: Drugs Drug Resistance. 14, 249–256.

25. Wheatley, N. C., Andrews, K. T., Tran, T. L., Lucke, A. J., Reid, R. C. & Fairlie, D. P. (2010) Antimalarial histone deacetylase inhibitors containing cinnamate or NSAID components, Bioorganic Medicinal Chemistry Letters. 20, 7080–7084.

26. Mok, S., Imwong, M., Mackinnon, M. J., Sim, J., Ramadoss, R., Yi, P., Mayxay, M., Chotivanich, K., Liong, K.-Y. & Russell, B. (2011) Artemisinin resistance in Plasmodium falciparum is associated with an altered temporal pattern of transcription, BMC Genomics. 12, 1–14.

27. Dovey, O. M., Foster, C. T. & Cowley, S. M. (2010) Histone deacetylase 1 (HDAC1), but not HDAC2, controls embryonic stem cell differentiation, Proceedings of the National Academy of Sciences. 107, 8242–8247.

28. Kulka, L. A. M., Fangmann, P.-V., Panfilova, D. & Olzscha, H. (2020) Impact of HDAC inhibitors on protein quality control systems: consequences for precision medicine in malignant disease, Frontiers in Cell Developmental Biology. 8, 425.

29. Milutinovic, S., Zhuang, Q. & Szyf, M. (2002) Proliferating cell nuclear antigen associates with histone deacetylase activity, integrating DNA replication and chromatin modification, Journal of Biological Chemistry. 277, 20974–20978.

30. Oh, M., Choi, I.-K. & Kwon, H. J. (2008) Inhibition of histone deacetylase1 induces autophagy, Biochemical Biophysical Research Communications. 369, 1179–1183.

31. Thurn, K. T., Thomas, S., Raha, P., Qureshi, I. & Munster, P. N. (2013) Histone deacetylase regulation of ATM-mediated DNA damage signaling, Molecular Cancer Therapeutics. 12, 2078–2087.

32. Wilting, R. H., Yanover, E., Heideman, M. R., Jacobs, H., Horner, J., Van Der Torre, J., DePinho, R. A. & Dannenberg, J. H. (2010) Overlapping functions of Hdac1 and Hdac2 in cell cycle regulation and haematopoiesis, The EMBO Journal. 29, 2586–2597.

33. Cacan, E., Ali, M. W., Boyd, N. H., Hooks, S. B. & Greer, S. F. (2014) Inhibition of HDAC1 and DNMT1 modulate RGS10 expression and decrease ovarian cancer chemoresistance, PLoS One. 9, e87455.

34. Huang, Z., Li, R., Tang, T., Ling, D., Wang, M., Xu, D., Sun, M., Zheng, L., Zhu, F. & Min, H. (2020) A novel multistage antiplasmodial inhibitor targeting Plasmodium falciparum histone deacetylase 1, Cell Discovery. 6, 1–15.

35. Mukherjee, P., Pradhan, A., Shah, F., Tekwani, B. L. & Avery, M. A. (2008) Structural insights into the Plasmodium falciparum histone deacetylase 1 (PfHDAC-1): A novel target for the development of antimalarial therapy, Bioorganic Medicinal Chemistry. 16, 5254–5265.

36. Patel, V., Mazitschek, R., Coleman, B., Nguyen, C., Urgaonkar, S., Cortese, J., Barker Jr, R. H., Greenberg, E., Tang, W. & Bradner, J. E. (2009) Identification and characterization of small molecule inhibitors of a class I histone deacetylase from Plasmodium falciparum, Journal of Medicinal Chemistry. 52, 2185–2187.

37. Pflum, M. K. H., Tong, J. K., Lane, W. S. & Schreiber, S. L. (2001) Histone deacetylase 1 phosphorylation promotes enzymatic activity and complex formation, Journal of Biological Chemistry. 276, 47733–47741.

38. Pease, B. N., Huttlin, E. L., Jedrychowski, M. P., Talevich, E., Harmon, J., Dillman, T., Kannan, N., Doerig, C., Chakrabarti, R. & Gygi, S. P. (2013) Global analysis of protein expression and phosphorylation of three stages of Plasmodium falciparum intraerythrocytic development, Journal of Proteome Research. 12, 4028–4045.

39. Treeck, M., Sanders, J. L., Elias, J. E. & Boothroyd, J. C. (2011) The phosphoproteomes of Plasmodium falciparum and Toxoplasma gondii reveal unusual adaptations within and beyond the parasites’ boundaries, Cell Host Microbe. 10, 410–419.

40. Blom, N., Gammeltoft, S. & Brunak, S. (1999) Sequence and structure-based prediction of eukaryotic protein phosphorylation sites, Journal of Molecular Biology. 294, 1351–1362.

41. Birnbaum, J., Flemming, S., Reichard, N., Soares, A. B., Mesén-Ramírez, P., Jonscher, E., Bergmann, B. & Spielmann, T. (2017) A genetic system to study Plasmodium falciparum protein function, Nature Methods. 14, 450–456.

42. Telles, E. & Seto, E. (2012) Modulation of cell cycle regulators by HDACs, Frontiers in Bioscience. 4, 831.

43. Zhu, L., Tripathi, J., Rocamora, F. M., Miotto, O., van der Pluijm, R., Voss, T. S., Mok, S., Kwiatkowski, D. P., Nosten, F. & Day, N. P. (2018) The origins of malaria artemisinin resistance defined by a genetic and transcriptomic background, Nature Communications. 9, 1–13.

44. Hott, A., Casandra, D., Sparks, K. N., Morton, L. C., Castanares, G.-G., Rutter, A. & Kyle, D. E. (2015) Artemisinin-resistant Plasmodium falciparum parasites exhibit altered patterns of development in infected erythrocytes, Antimicrobial Agents Chemotherapy. 59, 3156–3167.

45. Xiong, A., Prakash, P., Gao, X., Chew, M., Tay, I. J. J., Woodrow, C. J., Engelward, B. P., Han, J. & Preiser, P. R. (2020) K13-mediated reduced susceptibility to artemisinin in Plasmodium falciparum is overlaid on a trait of enhanced DNA damage repair, Cell Reports. 32, 107996.

46. Witkowski, B., Amaratunga, C., Khim, N., Sreng, S., Chim, P., Kim, S., Lim, P., Mao, S., Sopha, C. & Sam, B. (2013) Novel phenotypic assays for the detection of artemisinin-resistant Plasmodium falciparum malaria in Cambodia: in-vitro and ex-vivo drug-response studies, The Lancet Infectious Diseases. 13, 1043–1049.

47. Ismail, H. M., Barton, V., Phanchana, M., Charoensutthivarakul, S., Wong, M. H., Hemingway, J., Biagini, G. A., O’Neill, P. M. & Ward, S. A. (2016) Artemisinin activity-based probes identify multiple molecular targets within the asexual stage of the malaria parasites Plasmodium falciparum 3D7, Proceedings of the National Academy of Sciences. 113, 2080–2085.

48. Wang, J., Zhang, C.-J., Chia, W. N., Loh, C. C., Li, Z., Lee, Y. M., He, Y., Yuan, L.-X., Lim, T. K. & Liu, M. (2015) Haem-activated promiscuous targeting of artemisinin in Plasmodium falciparum, Nature Communications. 6, 1–11.

49. Rawat, M., Kanyal, A., Sahasrabudhe, A., Vembar, S. S., Lopez-Rubio, J.-J. & Karmodiya, K. (2021) Histone acetyltransferase PfGCN5 regulates stress responsive and artemisinin resistance related genes in Plasmodium falciparum, Scientific Reports. 11, 1–13.

50. Zhu, L., van der Pluijm, R. W., Kucharski, M., Nayak, S., Tripathi, J., Nosten, F., Faiz, A., Amaratunga, C., Lek, D., Ashley, E. A., Smithuis, F., Phyo, A. P., Lin, K., Imwong, M., Mayxay, M., Dhorda, M., Chau, N. H., Thuy, N. N. T., Newton, P. N., Jittamala, P., Tripura, R., Pukrittayakamee, S., Peto, T. J., Miotto, O., Seidlein, L. v., Hien, T. T., Ginsburg, H., Day, N. P., White, N. J., Dondorp, A. M. & Bozdech, Z. (2021) The mechanism of artemisinin resistance of Plasmodium falciparum malaria parasites originates in their initial transcriptional response, Biorxiv, 2021.05.17.444396.

51. Kawai, H., Li, H., Avraham, S., Jiang, S. & Avraham, H. K. (2003) Overexpression of histone deacetylase HDAC1 modulates breast cancer progression by negative regulation of estrogen receptor α, International Journal of Cancer. 107, 353–358.

52. Lasonder, E., Green, J. L., Grainger, M., Langsley, G. & Holder, A. A. (2015) Extensive differential protein phosphorylation as intraerythrocytic Plasmodium falciparum schizonts develop into extracellular invasive merozoites, Proteomics. 15, 2716–2729.

53. Davies, H., Belda, H., Broncel, M., Ye, X., Bisson, C., Introini, V., Dorin-Semblat, D., Semblat, J.-P., Tibúrcio, M. & Gamain, B. (2020) An exported kinase family mediates species-specific erythrocyte remodelling and virulence in human malaria, Nature Microbiology. 5, 848–863.

54. Quinlan, A. R. & Hall, I. M. (2010) BEDTools: a flexible suite of utilities for comparing genomic features, Bioinformatics. 26, 841–842.

55. Thorvaldsdóttir, H., Robinson, J. T. & Mesirov, J. P. (2013) Integrative Genomics Viewer (IGV): high-performance genomics data visualization and exploration, Briefings in Bioinformatics. 14, 178–192.

56. Ramírez, F., Dündar, F., Diehl, S., Grüning, B. A. & Manke, T. (2014) deepTools: a flexible platform for exploring deep-sequencing data, Nucleic Acids Research. 42, W187–W191.

57. Aurrecoechea, C., Brestelli, J., Brunk, B. P., Dommer, J., Fischer, S., Gajria, B., Gao, X., Gingle, A., Grant, G. & Harb, O. S. (2009) PlasmoDB: a functional genomic database for malaria parasites, Nucleic Acids Research. 37, D539–D543.

