## Supplemental Figure 1 for "PfHDAC1 is an essential regulator of parasite asexual growth with its altered genomic occupancy and activity associated with artemisinin drug resistance in *Plasmodium falciparum*"

(A)

```
>PF3D7_0925700  449 amino acids
#
# netphos-3.1b prediction results
#
# Sequence                    # x   Context     Score   Kinase    Answer
# -------------------------------------------------------------------
# PF3D7_0925700             391 S   DFFNSDIDD   0.645   CKII       YES
# PF3D7_0925700             397 S   IDDESDKNQ   0.632   CKII       YES
# PF3D7_0925700             440 S   FFDLSDRDQ   0.544   CKII       YES
```

(B)

```
    MSNRKKVAYFHDPDIGSYYYGAGHPMKPQRIRMTHSLIVSYNLYKYMEVY   #    50
    RPHKSDVNELTLFHDYEYIDFLSSISLENYREFTYQLKRFNVGEATDCPV   #   100
    FDGLFQFQQSCAGASIDGASKLNHHCADICVNWSGGLHHAKMSEASGFCY   #   150
    INDIVLGILELLKYHARVMYIDIDVHHGDGVEEAFYVTHRVMTVSFHKFG   #   200
    DYFPGTGDITDVGVNHGKYYSVNVPLNDGMTDDAFVDLFKVVIDKCVQTY   #   250
    RPGAIIIQCGADSLTGDRLGRFNLTIKGHARCVEHVRSYNIPLLVLGGGG   #   300
    YTIRNVSRCWAYETGVVLNKHHEMPDQISLNDYYDYYAPDFQLHLQPSNI   #   350
    PNYNSPEHLSRIKMKIAENLRHIEHAPGVQFSYVPPDFFNSDIDDESDKN   #   400
    QYELKDDSGGGRAPGTRAKEHSTTHHLRRKNYDDDFFDLSDRDQSIVPY    #   450
%1  .S..............SY...............T.S...S.....Y....   #    50
%1  ....S.....T......Y.....S.S........Y..........T....   #   100
%1  ...................S......................S......Y   #   150
%1  ...................Y...............Y.T....T.S.....   #   200
%1  .....T...T........YY..........T.................T.   #   250
%1  ............S...........T.............Y...........   #   300
%1  YT....S....Y.T..............S...YY.YY..........S..   #   350
%1  ....S....S..............................S.....S...   #   400
%1  .Y.....S.......T.....S.................S....S....
```

(C)

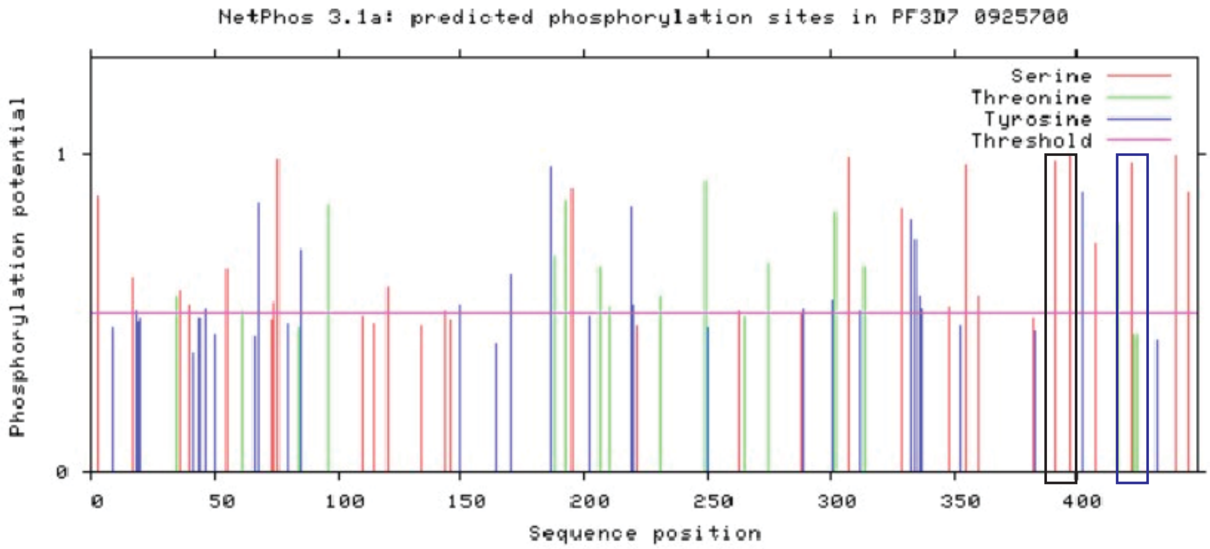
