## Supplementary figures and images for "PfHDAC1 is an essential regulator of parasite asexual growth with its altered genomic occupancy and activity associated with artemisinin drug resistance in *Plasmodium falciparum*"

### Supplemental Figure 2

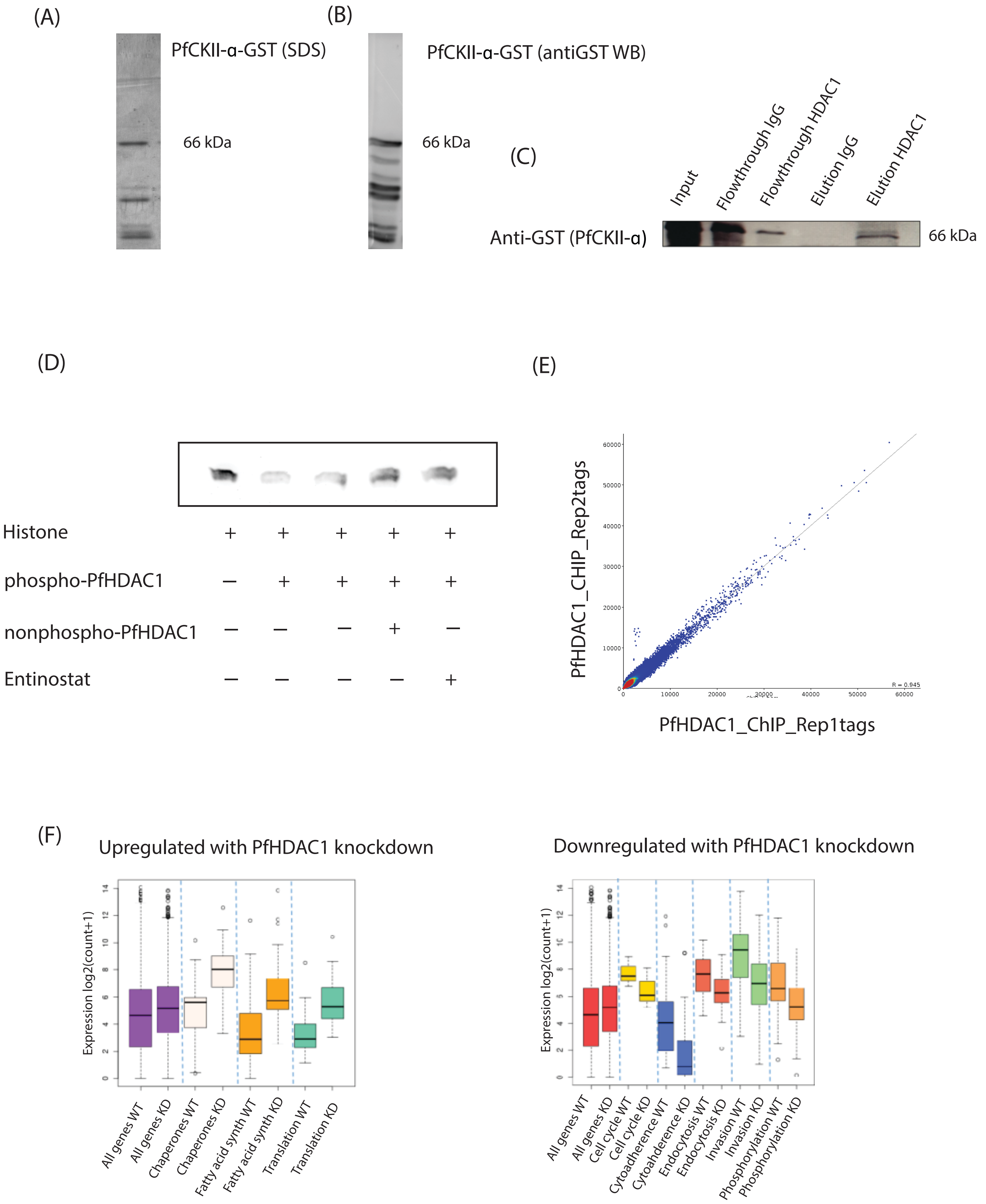

### Supplemental Figure 3

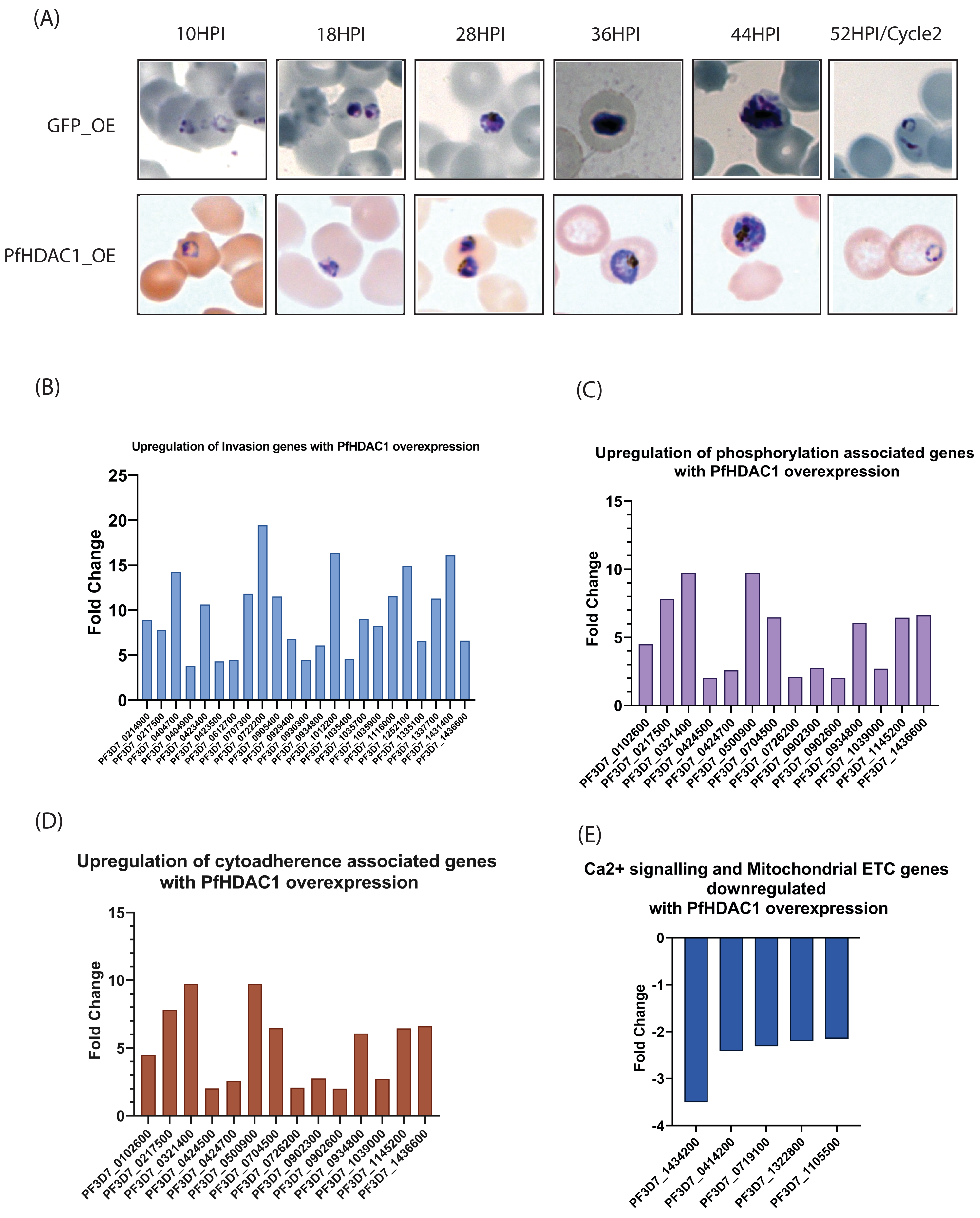

### Supplemental Figure 4

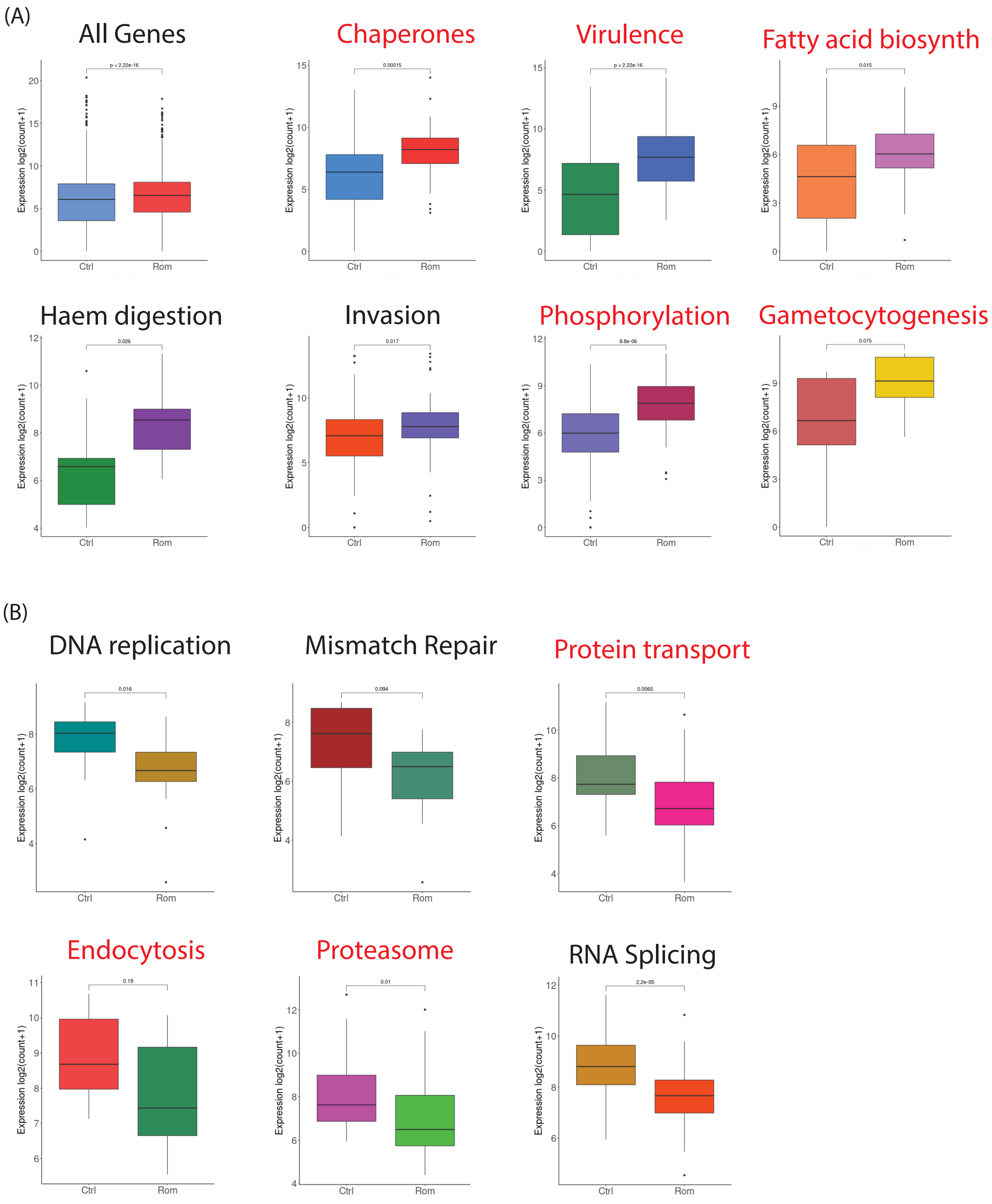

### Supplemental Figure 5

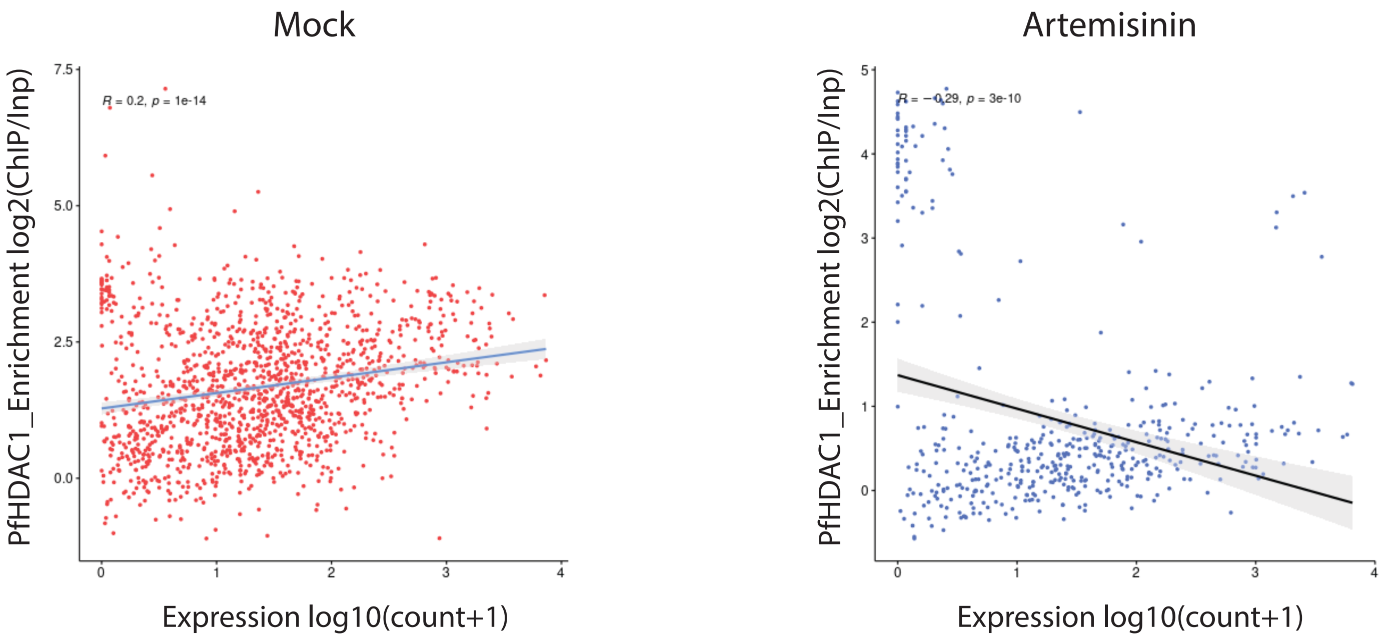
